# F-CPI: Prediction of activity changes induced by fluorine substitution using multimodal deep learning

**DOI:** 10.1101/2024.07.17.602844

**Authors:** Qian Zhang, Wenhai Yin, Xinyao Chen, Aimin Zhou, Guixu Zhang, Zhi Zhao, Zhiqiang Li, Yan Zhang, Jingshan Shen, Weiliang Zhu, Xiangrui Jiang, Zhijian Xu

## Abstract

There are a large number of fluorine (F)-containing compounds in approved drugs, and F substitution is a common method in drug discovery and development. However, F is difficult to form traditional hydrogen bonds and typical halogen bonds. As a result, accurate prediction of the activity after F substitution is still impossible using traditional drug design methods, whereas artificial intelligence driven activity prediction might offer a solution. Although more and more machine learning and deep learning models are being applied, there is currently no model specifically designed to study the effect of F on bioactivities. In this study, we developed a specialized deep learning model, F-CPI, to predict the effect of introducing F on drug activity, and tested its performance on a carefully constructed dataset. Comparison with traditional machine learning models and popular CPI task models demonstrated the superiority and necessity of F-CPI, achieving an accuracy of approximately 89% and a precision of approximately 67%. In the end, we utilized F-CPI for the structural optimization of hit compounds against SARS-CoV-2 3CL^pro^. Impressively, in one case, the introduction of only one F atom resulted in a more than 100-fold increase in activity (IC_50_: 22.99 nM vs. 28190 nM). Therefore, we believe that F-CPI is a helpful and effective tool in the context of drug discovery and design.

## 1. Introduction

### Fluorine (F) is of great importance in drug discovery and development

Fluorine (F) has a wide range of applications in pharmaceutical chemistry [1-6]. Due to its unique properties such as small atomic radius (1.47 Å) and strong electronegativity, F is commonly used to replace -H or -OH to improve various physicochemical properties of drug molecules [7, 8]. Since the first F-substituted drug, fluhydrocortisone, was launched in 1955, the introduction of F into drug molecules has been widely used and has shown a growing trend. For example, among the top 10 best-selling small molecule drugs in 2022, 5 contain F (Figure 1). Therefore, introducing F is of great importance in lead optimization.

**Figure 1.**
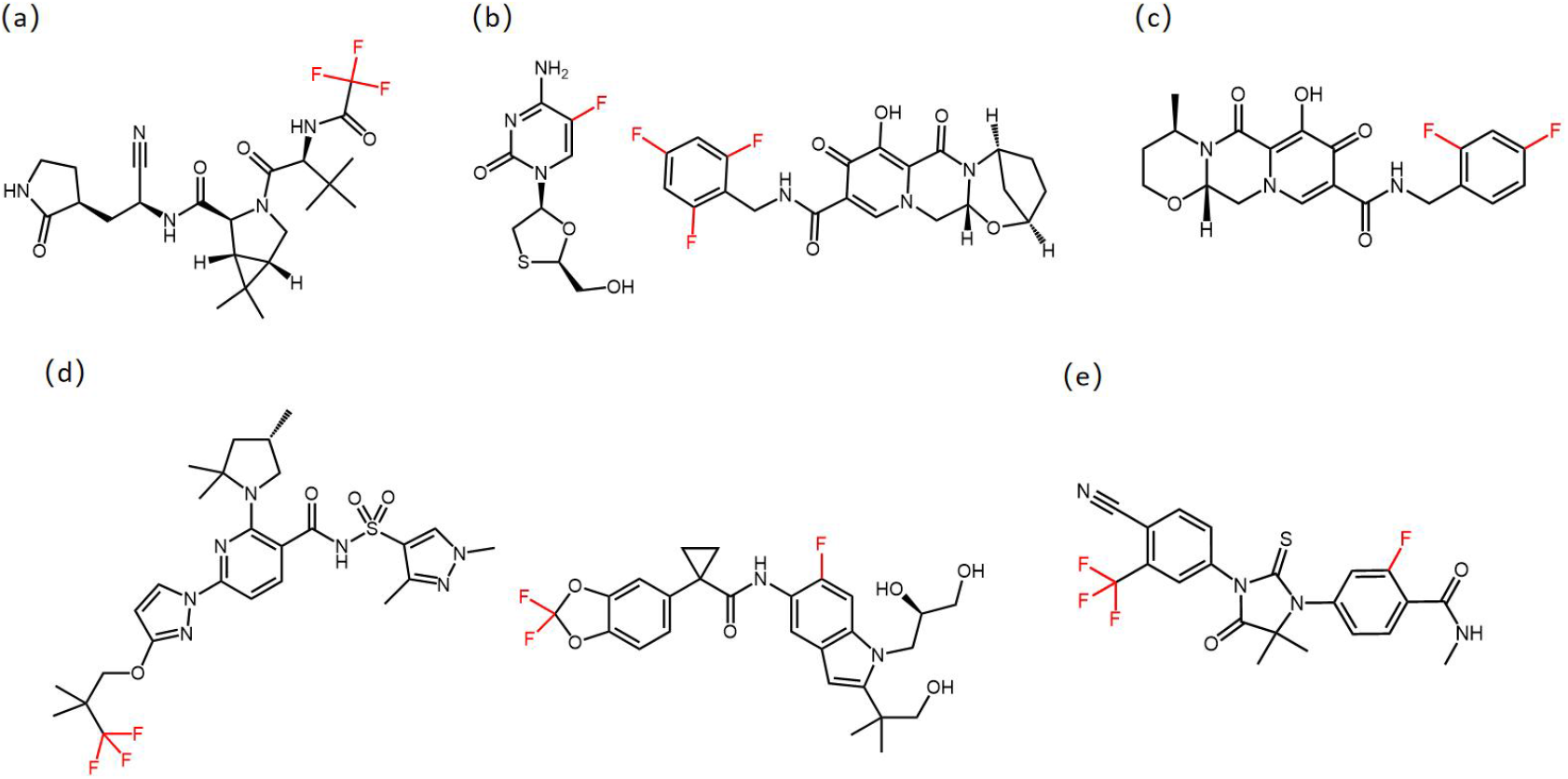
5 drugs containing F among the 10 best-selling small molecule drugs of 2022 (some drugs are in combination form and all components contain Fluorine). (a) Nirmatrelvir. (b) Emtricitabine (left) and Bictegravir (right). (c) Dolutegravir. (d) Elexacaftor (left) and Tezacaftor (right). (e) Enzalulutamide.

### F has a significant impact on the physicochemical properties of drug molecules

The introduction of F will change the charge distribution of drug molecules, significantly affecting the acidity and alkalinity of its neighboring groups, thereby affecting the pKa value of the compound. The introduction of F also enhances the lipophilicity of drug molecules, thereby enhancing their cell membrane permeability. Mono-F substitution typically exhibits weak hydrophobicity, while di-F substitutions exhibits strong hydrophobicity due to addition effect [9, 10]. In addition, C-F bonds are short and difficult to polarize, making them less prone to breakage. Introducing F at easily metabolizable sites can effectively improve the metabolic properties of compounds.

### F substitution can affect the activity of compounds

F substitution can enhance the binding affinity between compounds and targets (Figure 2). Our previous study indicated that 9.19% of compounds can increase their activity by at least an order of magnitude after replacing -CH_3_ with -CF_3_ [11]. F is difficult to form traditional hydrogen bonds and typical halogen bonds. As a result, accurate prediction of the activity after F substitution is still impossible using traditional drug design methods, while artificial intelligence driven activity prediction might become a solution.

**Figure 2.**
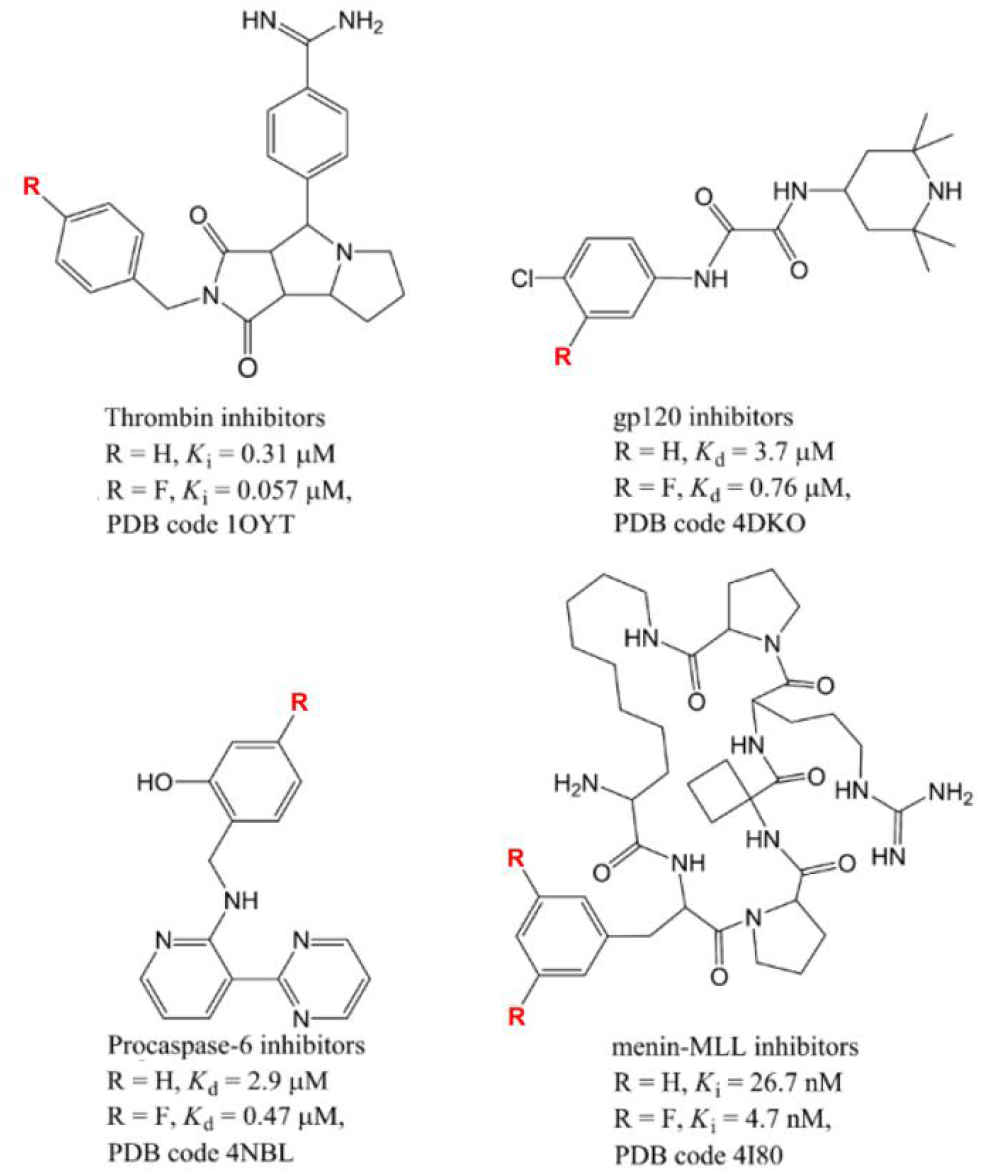
F substitution increases the binding affinity between compounds and target proteins. R represents the substituent group.

Compound-Protein Interaction by artificial intelligence to predict the activities of the compounds. Identifying compound-protein interactions (CPI) plays an important role in predicting the activity of compounds. Traditional methods for solving CPI tasks mainly rely on virtual screening based on physical modeling of the structures of compounds and proteins [12, 13]. These methods have been widely applied over the past few decades, but are often limited due to the difficulty in obtaining the 3D structure of proteins and the small size of molecular datasets. To address this issue, a large number of machine learning methods [14, 15] have emerged in recent years, such as Support Vector Machines (SVM) and Random Forests (RF). These methods do not rely on the 3D structure of proteins; thus, they have stronger generality and have achieved certain success in CPI tasks. With the rise of deep learning and the development of Graphics Processing Units (GPUs), more choices are available when faced with CPI tasks, from Multi-Layer Perceptron (MLP) [16], Recurrent Neural Networks (RNN) [17, 18], Convolutional Neural Networks (CNN) [19, 20], to the recently proposed Transformer with strong parallel capabilities [21-23], and Graph Neural Networks (GNN) [24, 25] that can better utilize molecular graph features. At the same time, traditional work models the CPI task as a binary classification problem, predicting whether a drug can bind a specific protein or not. Some new work [26] models the CPI task as a regression problem, attempting to predict the strength of the binding affinity.

The Effect of F Substitution on Drug Molecules. Identifying compound-protein interaction after F substitution (CPI-FS), is a subset of CPI tasks. In CPI-FS, F substitution refers to replacing a hydrogen atom in the original compound with a fluorine atom. The direct solution is to predict the binding affinity of compounds before and after F substitution, and calculate the activity changes. However, the activity cliff problem greatly increases the difficulty of model prediction[27]. The main challenges currently faced by this task are: (1) No publicly available dataset; (2) On this issue, the potential of deep learning has not been fully realized. In response to the above challenges, we have compiled a dataset, and proposed a general paradigm called Fluoro-substitution Compound-Protein Interaction Model (F-CPI) for solving CPI-FS tasks and introduce several prevailing deep learning methods. The best-performing model achieves approximately 67% precision, 89% accuracy, and 43% recall. In the end, we applied F-CPI to the structural optimization of hit compounds against SARS-CoV-2 3CL_pro_. Impressively, in one case, the introduction of only one F atom, the activity increased by more than two orders of magnitude (IC_50_: 22.99 nM vs. 28190 nM).

## 2. Materials and methods

### 2.1 Data Set of –F/-H Compounds

For comparing the effect of the F substitution on bioactivity, a data set was curated from ChEMBL, which is composed of matched molecular pair with bioactivity (one molecule with –F and the other with -H, while the rest of the paired molecular structures are exactly the same). The activities (K_i_, K_d_, IC_50_, EC_50_) are converted into negative logarithm units. The average will be used if multiple experiments are available for a compound against a target.

### 2.2 Overview

We consider CPI-FS tasks as binary classification problems based on compound and protein data, as shown in Figure 3. F-CPI aims to learn a mapping function that takes the protein features, original compound features, and F-substituted compound features as inputs and provides a binary prediction of 0 or 1, indicating whether there is a significant change in drug activity against the protein after F-substitution (i.e. ΔpKi >= 0.5).

**Figure 3.**
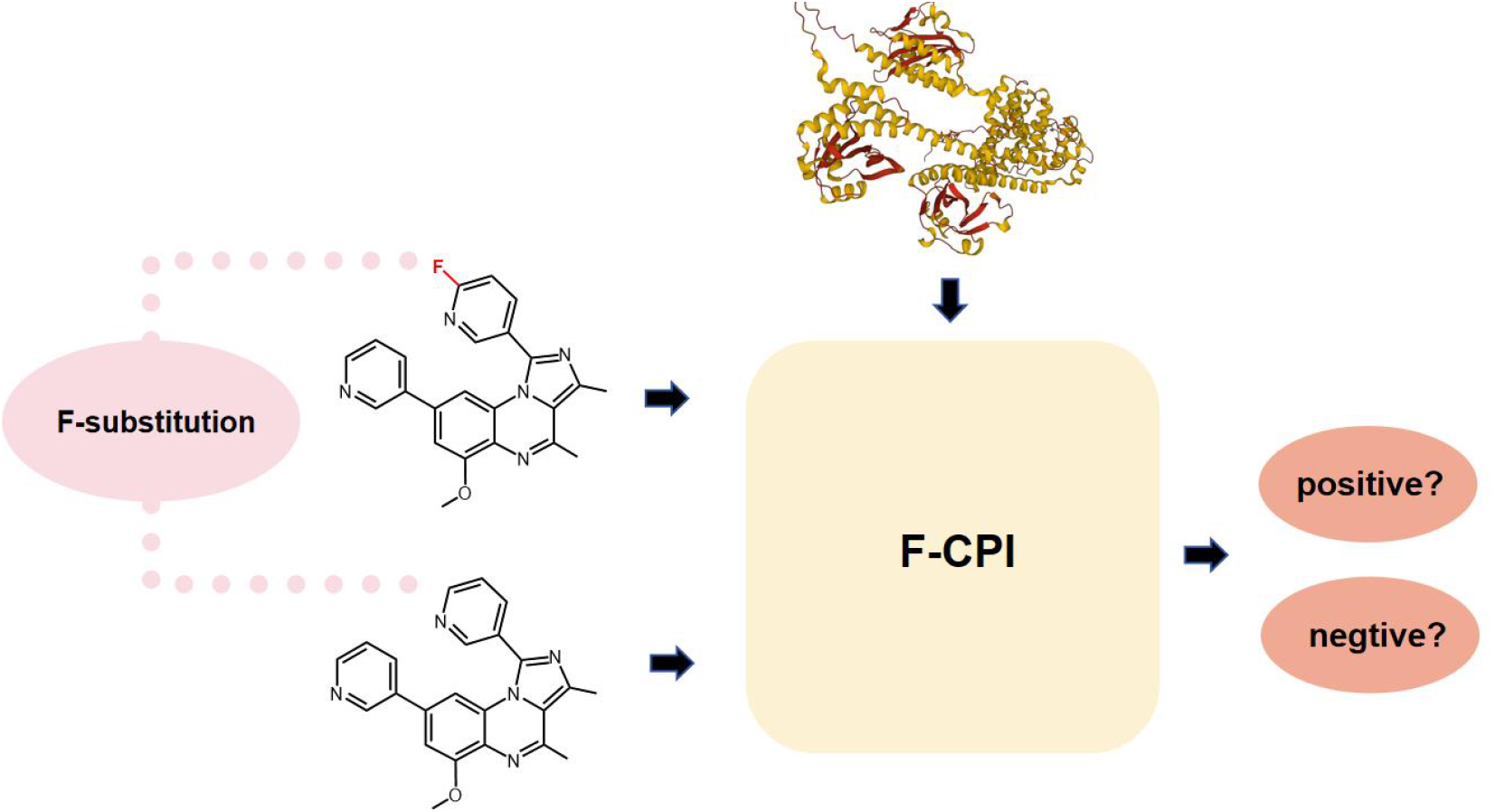
Identifying compound-protein interaction after F-substitution as binary classification problems.

We propose a universal model pattern Fluoro-substitution Compound-Protein Interaction Model (F-CPI) to solve the CPI-FS problem, as shown in Figure 4. The model consists of two encoders and one interactive decoding module. The encoders are used to extract high-dimensional hidden features of compounds and proteins, and the encoders are also responsible for fusing features from different modalities on the input side, such as molecular sequence features and molecular fingerprints. The fusion decoding module is used to simulate the interaction between compounds and proteins, and it fuses the features of compounds and proteins in an appropriate way to obtain the output result. It is worth mentioning that for the compound encoder, we adopted weight sharing technology to share the same encoder, which achieved better results in the experiment.

**Figure 4.**
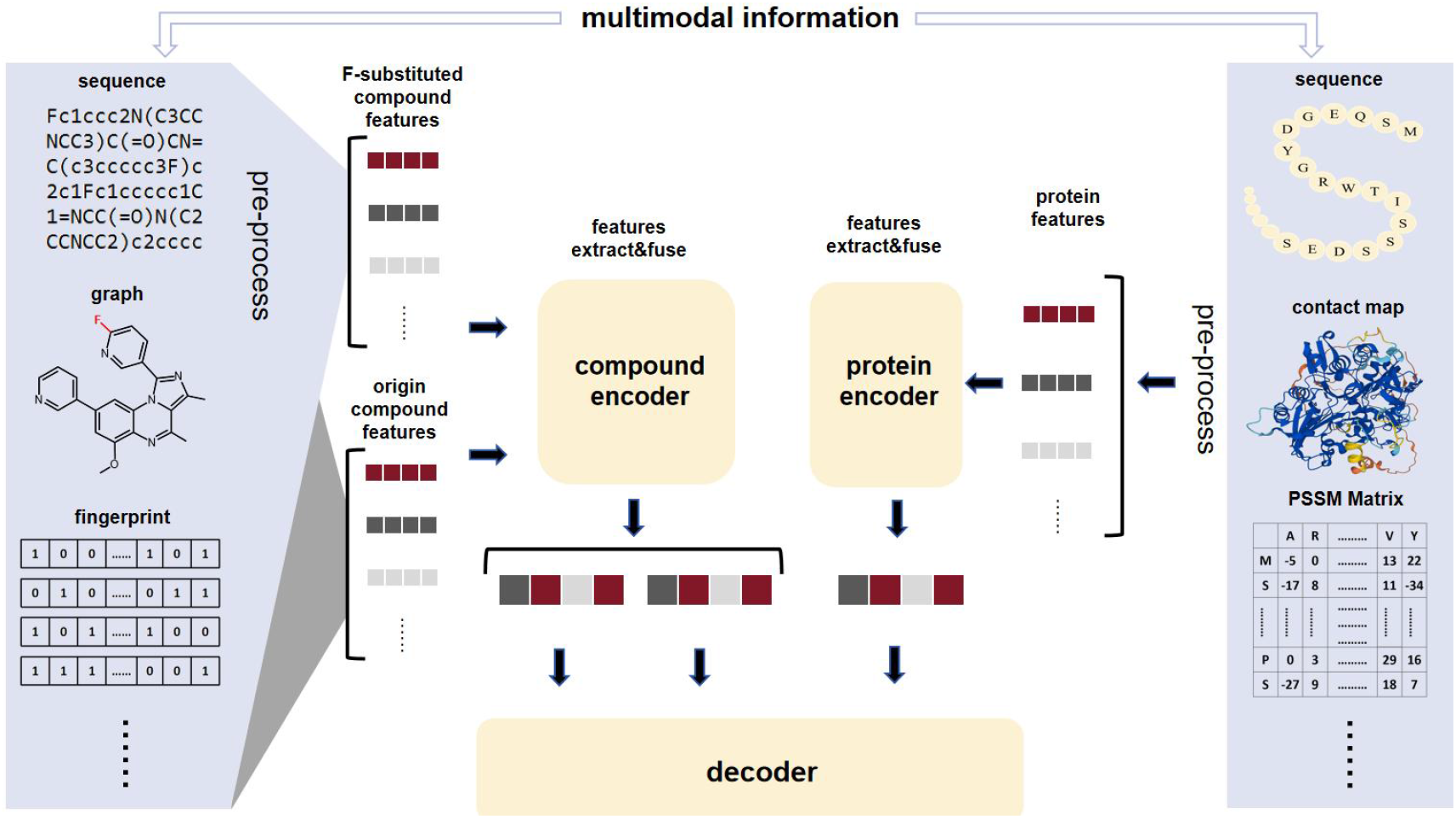
The overall framework of F-CPI. As shown on the left of the figure, the overlapping dialog box represents the parallel processing of F-substitution compound and the original compound in the preprocessing stage. In the compound encoding stage, we adopted weight sharing technology and used the same encoder for feature extraction and fusion.

### 2.3 Sequence-based Fluoro-substitution Compound-Protein Interaction Model (F-CPI (S))

We have designed a model called Sequence-based Fluoro-Substitution Compound-Protein Interaction Model (F-CPI (S)) based on sequence features. This model conforms to our proposed general pattern. We introduced sequence features of SMILES strings and amino acid sequences, and used sequence encoders for feature extraction. We use one layer for the compound encoder and three layers for the protein encoder with self-attention encoding layers. Finally, we obtain the higher-level feature maps 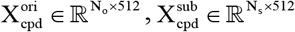 and 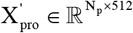,where N_o_, N_s_, N_p_ denote the original compound, F-substituted compound, and protein sequence lengths, respectively.

### Sequence encoder

For representing the SMILES sequence of a compound, we can treat each character as a word in a sentence. After passing through a learnable word embedding layer, the entire sequence is embedded into matrix 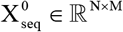, where N denotes the length of the sequence, M denotes the length of each word vector, and 0 represents the input of the first layer of the encoder. The Transformer [28] with self-attention mechanism can capture global information of the sequence and optimize local information. Due to its ability to fully utilize the parallel computing power of GPUs and reasonable structure, it has been widely applied and used in recent years in the field of drug chemistry for extracting sequence information using Transformer encoder. In this article, we also adopt a Transformer encoder based on the self-attention mechanism to extract sequence features of proteins or compounds. A classic self-attention mechanism is based on the following formulas:

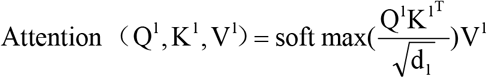

where Q, K, V represent query, key, value, obtained from 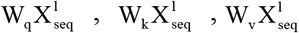 respectively. Where W ∈ ℝ^512×512^ denotes an unbiased linear layer, l∈(0, L), where L denotes the number of layers in the model. After one self-attention calculation on the sequence, it goes through a fully connected layer and dropout layer, followed by residual connection and layer normalization to obtain the output of the self-attention layer. To obtain the final result, we also need to pass the output of the self-attention layer through a Feedforward Neural Network (FFN):

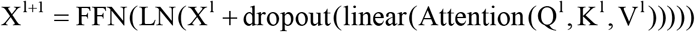

Where FFN () consists of two linear layers, an activation function, a dropout layer, a layer normalization layer, and a residual connection.

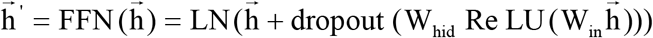

Where 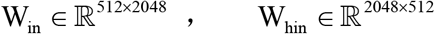 are set as default settings, LN() representing layer normalization, and the dropout is set to 0.1.

After repeating the above steps for several times, we finally obtain the representation matrix 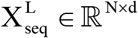 for sequence features.

### Sequence decoder

To better simulate the compound-protein interaction and comprehensively utilize the features of the entire sequence, we use the cross-attention mechanism for decoding. In the decoder layer, we use the feature vectors of two compounds as query, and the protein feature vectors as key and value for cross attention calculation

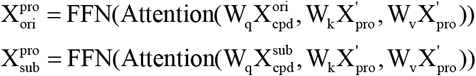

Where 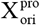 and 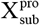 denotes the representation map of the interactions between the original compound and the F-substituted compound with proteins, respectively. Subsequently, perform forward and backward cross attention calculations on the two representation map, and use the difference as a representation of the entire interaction

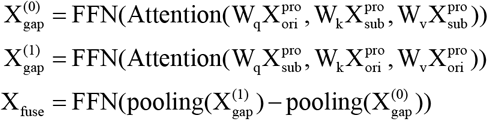

Where pooling() is the mean pooling function, which performs mean pooling operation on X _gap_ ∈ ℝ^L×512^ along the first dimension to obtain X _gap_ ‘ ∈ ℝ^1×512^.

### Focal-loss

In the constructed data set, there is a significant imbalance between positive and negative samples, as well as an imbalance in the difficulty level of sample prediction. To solve this problem, we adopt the focal loss method [29], which adjusts the contribution of each sample to the loss function to solve the problem of imbalanced positive and negative samples and imbalanced difficult and easy samples. For a general cross-entropy loss function:

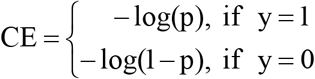

where y denotes the sample label and p denotes the probability value output by the model. Focal loss is proposed as follows:

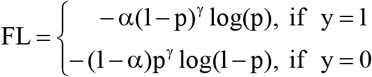

where *α* ∈(0,1) is used to adjust the contribution of positive and negative samples to the loss function, and *γ* is used to adjust the contribution of difficult and easy samples to the loss function. High confidence samples are considered easy samples and their weight is reduced, while low confidence samples are considered difficult samples and their weight is increased, aiming to make the model focus on difficult samples.

### Auxiliary tasks

To improve the performance of the model, we additionally design an auxiliary task module. The auxiliary task is a task set in addition to the main task, which is related to the main task. The auxiliary task guides the model’s learning direction during the gradient descent process by setting an additional loss function, thereby enhancing the model’s learning ability and helping the model achieve better performance on the main task [6]. We choose to predict the reaction activity values of the original compound and the F-substituted compound to the same protein as our sub-task. Due to the different indicator type of the compounds in the dataset, i.e., K_i_, IC_50_, EC_50_, etc., we need to introduce seven types of drug indicator features and encode them into one-hot vectors for word embedding, resulting in X _ind_ ∈ℝ^1×128^. For the input of the encoder, the original compound 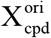, the F-substituted compound

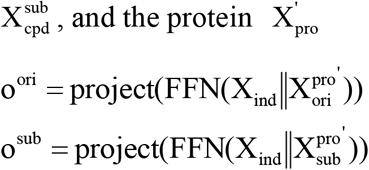

Where ‖ represents concatenation operation, 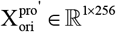 and 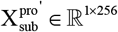 are obtained from 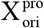 and 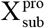 through a linear layer (512× 256) to ensure that the size of the output of the encoder is fixed. o^ori^ and o^sub^ denote the predicted drug activity values of the model for the original compound and F-substituted compound, respectively.

We choose the MSE function as the loss function for predicting activity values

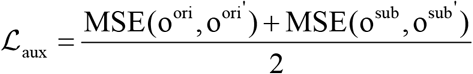

Where 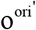 and 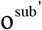 denote the ground truth of the drug activity values of the original compound and F-substituted compound, respectively. The overall loss of our final model is:

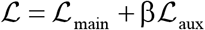

In the ablation experiment, we will discuss the selection of β.

### Pre-trained word embedding

Given a sequence, a common approach is to use a trainable simple word embedding layer during model training to embed tokens into a suitable vector space. In order to better explore the inherent high-dimensional information of proteins and compounds and further improve model performance, we introduce existing sequence-based pre-training models chemBERTa [30] and TAPE [31] to replace our word embedding layer. chemBERTa is a molecule pre-training model based on BERT [32], chemBERTa provides multiple pre-trained models, and the model used in this article is trained on PubChem77M (77 million) dataset. It trained with hidden size 384, dropout rate 0.1, the number of multi-head of attention is 12, and has 3 hidden layers. TAPE is a protein pre-training model trained on the Pfam dataset (31 million). the model used in this article based on BERT with hidden size 768, dropout rate 0.1, the number of multi-head of attention is 12, and has 12 hidden layers. we adopt the pre-training word embedding strategy by treating the entire pre-trained model as the word embedding layer of F-CPI(S) and further training it within the model. Tape and chemBERTa provide different embedding methods, such as full sequence embedding, pooling embedding, and mean embedding. For our proposed model F-CPI(S), we adopt the full sequence embedding. Two pre-trained word embedding layers take protein and compound sequences as inputs and output 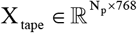 and 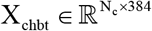 as word embedding representations, respectively. Where N_p_ denotes the length of the protein sequence, N_c_ denotes the length of the compound sequence. Finally, we use a linear layer to map the lengths of the compound output and protein output to a unified length of 512. We compared the effects of two word embedding methods in the result section, named F-CPI(S) and F-CPI(S-emb) respectively.

### 2.4 Graph-based fluoro-substitution compound-protein interaction model (F-CPI (G))

Due to the high degree of fit between compound structure and graph structure, Graph neural network (GNN) has been used for feature extraction of compounds in an increasing number of research works and has achieved certain success. Meanwhile, with the gradual improvement of protein 3D structural databases, in order to better utilize protein 3D structural information, some studies [33] attempt to convert structural information into protein contact maps, indirectly introducing structural features into the model through GNN.

Inspired by this, we designed a graph-based model called Graph-based Fluoro-Substitution Compound-Protein Interaction Model (F-CPI (G)).

#### Compound input

For compounds, we consider the atoms in a molecule as nodes, and the bonds between atoms as edges. We convert the original SMILES strings into node set V and edge set E using the RDKit [34] library. V contains all the atoms in the molecule, and we process atom features as follows:

(1) Atom types,

(2) Hybridization types,

(3) Atom degrees indicating the number of bonded neighboring atoms,

(4) Whether an atom is part of an aromatic ring or not,

(5) Formal charge,

(6) Chiral type,

(7) Implicit valence for atoms.

For the above atom features, they can be considered as 7 one-hot vectors. We introduce seven word embedding layers to individually embed these features, and then concatenate them into a vector with a length of 512 as the initial atom feature 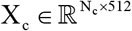, where N_c_ represents the number of atoms in the compound. The edge set of the graph is represented by the adjacency matrix A_c_ ∈ ℝ^N×N^, which indicates the connectivity information between nodes (atoms) in the molecule. a_i, j_ = 1 represents the existence of a chemical bond between the i-th and j-th atoms.

#### Protein input

Considering amino acids as nodes and embed them by their types and obtain 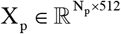. N_p_ represents the number of amino acids in the protein. To represent the connectivity information between nodes, we introduce a protein contact map. For a protein with a length of L, its contact map is a square matrix C = {c_p,q_}.

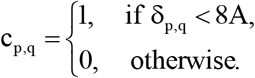

Where δ_p,q_ denotes the Euclidean distance between the two C_*α*_ atoms of the p-th and q-th residues. Generally, if δ_p,q_ is less than 8Å, two residues are defined as in contact. We use the contact map as the adjacency matrix input to the GNN.

AlphaFold [35] can predict protein structures with atomic-level precision. We obtained most of the predicted 3D protein structures from the AlphaFold database (https://alphafold.ebi.ac.uk/). For a small number of samples not included in the AlphaFold database, we used the RoseTTA-fold [36] model to predict protein structures and submitted our requests through the online version of Robetta (https://robetta.bakerlab.org/).

#### Encoder using Graph Convolutional Networks(GCN)

We use GCN [37] as the encoder for proteins and compounds. For a given graph structure, the aggregation process is as follows:

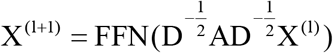

where D denotes the degree matrix of the adjacency matrix, and l denotes the GCN layer number.

#### Encoder using Graph Attention Networks(GAT)

As a comparison, we introduce GAT [38] as the encoder for proteins and compounds. The similarity between queries and keys is calculated using the dot product method. For a given graph structure:

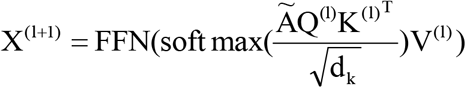

where 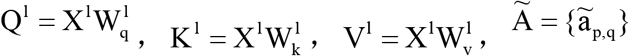

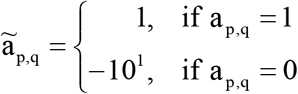

It is worth mentioning that in classical GAT, when a_p,q_ = 0, there is usually ã_p,q_ = −10^9^, which may lead to the model overly focusing on local information and ignoring global information. This article smoothed it to enable the model to have higher attention to amino acid residues that are close in distance while accessing global information.

#### Graph readout

We adopted a mean-weighted readout approach to obtain a high-dimensional representation of the entire graph structure. For the output of the last layer X^L^, which can be seen as a set of node features 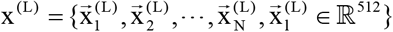, we have:

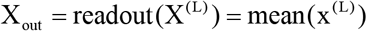

After applying the graph neural network to extract features from compounds and proteins, followed by readout, we finally obtain 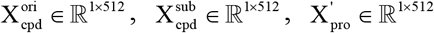.

#### Decoder

In the decoding stage, we adopt a decoding method based on concatenation fusion. We concatenate the three outputs 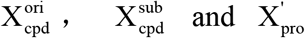 of the encoders along the last dimension, and then pass them through a FFN, followed by a linear layer for dimension reduction, resulting in X_fuse_∈ℝ^1×512^. The formula is as follows:

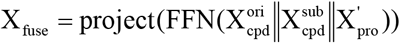

Where project () denotes a linear layer applied to(1536 × 512).

Finally, we use a classifier (linear layer) to obtain X_out_ ∈ℝ^1×2^ for binary classification loss calculation. F-CPI (G) also adopts focal loss function and auxiliary task method.

#### Auxiliary tasks

We designed auxiliary tasks suitable for F-CPI (G). Due to the different decoder structure from F-CPI (S), we adopted a concatenating approach to design the auxiliary task module

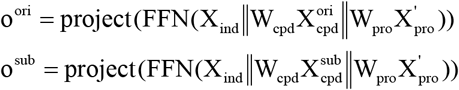

### 2.5 Multimodal Fluoro-Substitution Compound-Protein Interaction Model (F-CPI (M))

Furthermore, we propose a dedicated model called Multimodal Fluoro-Substitution Compound-Protein Interaction Model (F-CPI (M)) for predicting the activity of F-substituted compounds. Under the general pattern mentioned earlier, we combine multi-modal inputs to ensure satisfactory performance of the model.

#### Multimodal compound Encoder

In the compound encoder, we handle both sequence features and Morgan fingerprint features of the compounds in parallel.

For sequence features, we embed them into a matrix 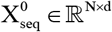 using a custom vocabulary and a learnable word embedding layer. N denotes the length of the SMILES string, d denotes the feature length (usually set as 512). Then a layer of self-attention based transformer encoder was used, which is the same as the F-CPI (S) model. Differently, F-CPI (M) sets a mean pooling layer at the end of the encoder, resulting in 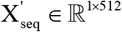.

For Morgan fingerprint features, we extract features using a linear layer and a FFN to obtain 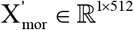. Then, we concatenate 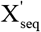 and 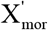, and pass them through a FFN and a linear layer for fusion and dimension reduction, resulting in 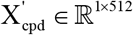. The parameters of the original compound encoder and F-substituted compound encoder are shared.

#### Morgan fingerprint

Morgan fingerprint, which is a characteristic representation method used in chemoinformatics to describe molecules [39], it takes into account the topological distance between atoms and the connectivity degree. It generates a binary vector representation by iteratively combining each atom with its neighboring atoms, effectively capturing the local information of molecules. Due to its ability to specify the length, it has been widely used in machine learning. In this study, we introduce Morgan fingerprint as a chemical descriptor for drug molecules in machine learning-based methods, enabling the model to have predictive capability. Furthermore, in F-CPI (M), we fuse the Morgan fingerprint features with the sequence features of compounds, achieving improved performance. The radius of Morgan fingerprint in this study is set to 4.

#### Multimodal Protein Encoder

In the protein encoder, we integrate a sequence encoder based on pre-trained word embedding methods and a PSSM matrix obtained from sequence alignment. We adopt the TAPE model as the pre-training word embedding layer to embed the amino acid sequences into vector representations. Unlike F-CPI (S), we chose the mean-pooling word embedding method provided by TAPE to obtain X _tape_ ∈ ℝ^1×768^, followed by a linear layer and FFN to finally obtain 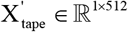.

Additionally, we incorporate PSSM-400 [40] as a high-level protein feature, denoted as X _PSSM_ ∈ ℝ^1×400^. After passing through a linear layer and an FFN, we obtain 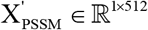.

Finally, we concatenate 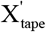 and 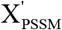, and fuse and reduce their dimensionality through a FFN and a linear layer, ultimately obtaining 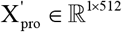 as the high-dimensional feature map of the protein.

#### PSSM-400

In previous studies, Position-Specific Scoring Matric (PSSM), which contain protein evolution information, have been widely used in predicting protein secondary structures. For a protein sequence of length N, the corresponding PSSM matrix is P∈ℝ^N×20^. Each entry represents the likelihood of an amino acid mutating into one of the 20 amino acids. In order to map the irregular matric into a fixed-length vector and adapt them to machine learning, normalizing the matrix values between 0 and 1 using the formula

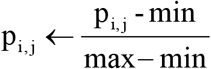

Where max and min represent the maximum and minimum values in PSSM, respectively. Then, sum the vectors corresponding to the same type of amino acid in the sequence, resulting in 20 vectors of length 20. These vectors are then concatenated to obtain a protein PSSM-400 vector X_PSSM_ ∈ ℝ^1×400^. Our PSSM matrices are obtained through the psi-blast alignment algorithm on the Swissport database, with an algorithm E-value of 0.01, 3 iterations, and default parameters for other settings.

#### Decoder

In the decoding stage, we use the same decoding method based on concatenation fusion as the F-CPI (G) model, along with focal loss function and auxiliary tasks method.

### 2.6 Traditional Machine Learning Algorithms

Traditional machine learning typically performs well on small sample datasets, while using naive machine learning in pharmaceutical chemistry often yields good results [41, 42]. To compare with our proposed model, we selected Support Vector Machine (SVM) and Random Forest (RF). For machine learning methods, we utilized the existing sklearn package. Since these methods require fixed-length input vectors, we used the Morgan fingerprint with a radius of 4 as the compound feature, resulting in a vector of length 384. For proteins, we used the PSSM-400 encoding method to represent the amino acid sequence as a vector of length 400. Finally, we concatenated the two compound feature vectors and the protein feature vector obtained above, resulting in a final input vector of length 1168 for the machine learning model. For each approach, the model hyperparameters were optimized as follows: (a) SVM, optimization of the kernel coefficient (γ) and regularization parameter (C), γ = [0.00001, 0.0001, 0.001, 0.01, or 0.1] and C = [1, 10, 100, 1000];(b) RF, number of decision trees (t), t = [100, 500, 1000,10000].

## 3. Results

This article proposes a general pattern for predicting the activity changes of compounds after F-substitution, and designs three distinctive deep learning models, while conducting reasonable feature engineering to introduce machine learning methods. In this section, we compared the performance of various methods using different evaluation metrics on a specially constructed dataset. Finally, taking the best performing F-CPI (M) as an example, ablation experiments were conducted to investigate the contribution of each module of the model to the task.

## Datasets

A total of 222,336 pairs of compound-protein reactions that can be paired were collected through online databases and previous papers, resulting in 111,168 samples.

Each sample consists of

(1) original and F-substituted drug compound SMILES strings

(2) protein amino acid sequences

(3) types of drug activity indicator

(4) logarithmic activity values for the reactions of each compound with the protein

(5) the difference between the two values

The dataset includes 2,503 proteins, 73,787 compounds, and 7 types of activity measurements. We divided the training set, validation set, and test set in a 9:1:1 ratio. Activity cliff refers to a situation where two structurally similar molecules exhibit significantly different activity values. In general, it is considered an activity cliff when the difference in logarithmic activity value for two structurally similar molecules exceeds 2 (equivalent to a difference of 100-fold in activity). Inspired by this, we define a significant improvement as a difference in logarithmic activity of 0.5 or greater (approximately 3.16-fold increase) when introducing fluorine into drug molecules. Based on this criterion, we created 14,679 positive samples and 96,489 negative samples.

### Evaluation metrics

We use three common evaluation metrics to assess the results of binary classification predictions: accuracy, precision, and recall. They are defined as follows:

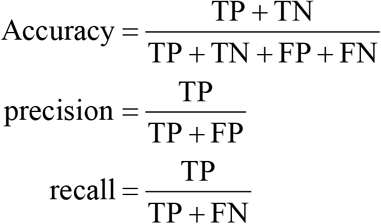

Here, TP, TN, FP, and FN represent true positives, true negatives, false positives, and false negatives, respectively.

### Setting

We conducted experiments using PyTorch toolkit in Python. We also utilized the powerful computing capabilities of NVIDIA RTX3090. The model was trained using the ADAM optimizer for 500 epochs, with a learning rate of 0.3 and a warm-up of 2000 steps. The batch size during training was set to 144, and the gradient accumulation size was set to 2. A dropout of 0.1 was used to mitigate overfitting, and early stop technique was employed to prevent overfitting, the strategy is to stop training when the accuracy of the model does not improve for 100 consecutive epochs on the validation set.

### 3.1 F-CPI outperforms other methods

Since there is no prior work on predicting changes in drug activity after F-substitution using machine learning or deep learning, we compared the performance of mainstream machine learning methods and deep learning techniques in this task as comprehensively as possible on the constructed dataset, as shown in Table 1.

**Table 1.** Comparison of performance between several machine learning and deep learning methods.

| Methods | ACC(%) | RECALL(%) | PRE(%) |
| --- | --- | --- | --- |
| SVM | 86.39 | 26.74 | 45.59 |
| RF | 87.39 | 22.75 | 59.31 |
| SVM(p-emb) | 87.17 | 38.44 | 53.87 |
| RF(p-emb) | 87.30 | 22.68 | 58.33 |
| F-CPI (GCN) | 86 | 27.17 | 47.31 |
| F-CPI (GAT) | 86.4 | 36.30 | 50.31 |
| F-CPI (S) | 87.46 | 22.68 | 60.16 |
| F-CPI (S-emb) | 88.23 | 23.12 | **70.25** |
| F-CPI (M) | **89.46** | **43.59** | 67.27 |

Note: p-emb represents pre-trained embedding, S-emb represents pre-trained embedding & sequence-base. Bold font indicates the best result.

F-CPI (M) has achieved excellent performance in accuracy and recall, while F-CPI (S-emb) has achieved excellent performance in precision. In specific application scenarios, pharmaceutical chemists need to conduct wet experiments on F-substituted compounds to verify the results, and also need to perform a series of synthesis steps to obtain the F-substituted molecules. Under the influence of experimental costs, we believe that the precision (In the F-substitution cases, the proportion of compounds with observed increased activity to compounds predicted with increased activity) of prediction is more important than the recall rate. Our goal is to achieve the highest possible accuracy while maintaining a certain recall rate. But compared to F-CPI (M), although F-CPI (S-emb) is 0.58% higher in precision, it is 1.33% and 20.32% lower in accuracy and recall, respectively, indicating that F-CPI (M) has better overall performance.

It is worth mentioning that the proportion of positive and negative samples in the test set is close to 1 to 7. In such a large proportion, the performance of the model can help pharmaceutical chemists screen out a large number of invalid substitutions and ensure that the results have a certain level of confidence, which has practical significance. Specific applications can be found in case studies.

For machine learning, good feature engineering has a significant impact on the model’s performance. Molecular fingerprinting is a widely used and verified feature engineering method in molecular machine learning, which even surpasses deep learning methods in certain specific tasks such as activity cliffs. However, there is no unified paradigm for protein feature engineering. Although we have tried to use more comprehensive PSSM matrices for processing, in order to further demonstrate the advantages of F-CPI, we replaced the proposed protein PSSM-400 feature vectors with pre-trained word embedding vectors based on the TAPE pre-training model as inputs for traditional machine learning methods, named p-emb. The results are shown in rows 3-4 of the table, and it can be seen that there is indeed an improvement in the results for SVM, but there is still a certain gap compared to F-CPI in various indicators.

For F-CPI (G), the results are shown in rows 5-6 of the table. GAT has improved accuracy by 1.21%, recall by 7.29%, and precision by 5.68% compared to GCN, which we attribute to the effectiveness of attention mechanism and attention mask smoothing operation. However, both methods do not have advantages compared to traditional machine learning methods.

For F-CPI (S), the introduction of pre-trained word embedding improved the accuracy of the model by 1.01%, recall by 2.06%, and precision by 11.48% (Table 2).

**Table 2.** Comparison of performance between several CPI models for regression on dataset filtered by IC_50_.

| Methods | ACC(%) | RECALL(%) | PRE(%) | MSE |
| --- | --- | --- | --- | --- |
| DeepDTA | 83.21 | 29.21 | 43.46 | 0.46 |
| GraphDTA (GIN) | 85.94 | 34.58 | 57.43 | 0.36 |
| GraphDTA (GAT) | 84.4 | 16.24 | 48.44 | 0.45 |
| GraphDTA (GAT&GCN) | 86.29 | 40.09 | 58.10 | **0.36** |
| GraphDTA (GCN) | 86.75 | 35.77 | 62.34 | 0.37 |
| F-CPI (M) | **89.65** | **44.71** | **79.16** | 0.42 |

Note: Bold font indicates the best result.

### 3.2 F-CPI outperforms other CPI models on dataset filtered by IC_50_

As mentioned earlier, there are some models for regression prediction of compound-protein binding affinity for traditional CPI tasks. To further evaluate the performance of F-CPI in predicting drug activity after F-substitution and demonstrate the necessity of an F-substitution specific model, we compared the performance of F-CPI and several other CPI models on regression and F-substitution tasks on a dataset filtered with IC_50_ as shown below.

Due to the traditional CPI model processing only one type of drug activity indicator, such as IC_50_, Ki or Kd [19, 24]. In order to better compare, we constructed a new dataset based on the original dataset filtered by IC_50_. It contains 1965 proteins, 42406 compounds, and a total of 48387 fluoride substituted control groups. In the end, we obtained a training set of size 39694, a validation set of size 4346, and a testing set of size 4347.

We introduced two popular CPI models DeepDTA [19] and GraphDTA [24] for comparison, and implement the following strategies to compare with F-CPI: (1) Extract all compound protein complexes from the training and validation sets, predict drug activity values, and select epoch and hyperparameters based on the performance of the validation set. (2) On the test set, a control group contains two pairs of compound-protein complex, then use the trained model to predict the activity values of the two pairs of complexes separately. The final prediction of this model is whether the improvement in activity after F-substitution is greater than 0.5 (logarithmic activity), and the accuracy, precision, and recall are calculated based on this. As shown in Table 2, F-CPI achieved the best performance in accuracy, precision, and recall. We also compared the auxiliary task MSE loss of the F-CPI model with other models, and the results showed that although predicting drug activity is a strategy for auxiliary its main tasks for F-CPI, it can still defeat some models that prioritize activity prediction as their main task. This means that F-CPI also has a good ability to handle traditional CPI problems and is proficient in F-substitution issues on this basis.

Compared with the suboptimal GraphDTA (GAT&GCN), the accuracy has increased by 3.36%, the recall has increased by 4.62%, and the precision has increased by 21.06%. The overall performance has significantly improved, which proves the necessity of a dedicated model for F-substitution problems.

## 4. Ablation study

In this section, we selected the F-CPI (M) with the best overall performance for ablation experiments to demonstrate the effectiveness of each module and study their contribution to the CPI-FS task.

### 4.1 Modules effect study

As shown in Table 3, in lines 1-2, the introduction of chemical prior features improved the predictive performance of F-CPI (M), indicating the effectiveness of multimodal fusion methods. Among them, the introduction of molecular Morgan fingerprint significantly improved the recall of the model by 13.55%, accuracy by 1.34% and precision by 4.39%, while the introduction of PSSM-400 had a smaller impact on performance. We attribute this to the complexity of proteins, which is difficult to be well expressed through manual prior design. As shown in the first and third rows of Table 1, the superiority of pre-trained embedding features compared to PSSM-400 also supports the point of view.

**Table 3.** Study of individual model components or method.

| Methods | ACC(%) | RECALL(%) | PRE(%) |
| --- | --- | --- | --- |
| w/o pssm-400 | 89.43 | **43.81** | 66.93 |
| w/o morgan | 88.21 | 28.06 | 65.35 |
| w/o focal loss | 89.17 | 40.21 | 66.83 |
| w/o aux-task | 87.99 | 32.84 | 60.68 |
| base | **89.46** | 43.59 | **67.27** |

Note: W/o morgan means that morgan fingerprint features are not fused in the encoder, while aux-task represents auxiliary task. Bold font indicates the best result.

As shown in the third row of Table 3, the introduction of focal loss improved the accuracy of the model by 0.47%, recall by 1.69% and precision by 2.69%. As shown in Tables 4, we further conducted parameter experiments on the coefficients α and γ of focal loss. With the decrease of α, the model achieved higher precision. This is because a smaller α encourages the model to make more conservative predictions for positive samples to improve the confidence of positive sample predictions, which is what we expect. However, when α is too small, the model becomes overly conservative, resulting in a sharp drop in recall while obtaining lower precision in 1.55%. This is because the model unreasonably amplifies the loss value of negative samples, resulting in an inability to learn the characteristics of positive samples well. With the increase of γ, the overall performance of the model improves until γ equals 5, indicating that focusing more on difficult-to-distinguish samples can enhance model performance. It is worth noting that when γ=0, that is, not distinguishing between easy and difficult samples, the predicted precision significantly improved, but recall was significantly reduced. In order to obtain relatively balanced results and higher accuracy, we did not use this parameter value.

**Table 4.** Performance of using different values of hyper-parameters for focal loss and auxiliary task.

| Coefficient | Value | ACC(%) | RECALL(%) | PRE(%) |
| --- | --- | --- | --- | --- |
| $\alpha$ | 0.3 | 87.39 | 15.76 | 64.65 |
|  | 0.4 | **89.46** | 43.59 | **67.27** |
|  | 0.5 | 89.32 | 44.92 | 65.59 |
|  | 0.6 | 89.05 | 46.02 | 63.32 |
|  | 0.7 | 88.91 | **47.72** | 61.89 |
| $\gamma$ | 0 | **89.57** | 35.13 | **74.65** |
|  | 1 | 89.24 | 41.09 | 66.91 |
|  | 3 | 89.3 | 42.86 | 66.44 |
|  | 5 | 89.46 | **43.59** | 67.27 |
| $\beta$ | 0 | 87.99 | 32.84 | 60.68 |
|  | 0.1 | 89.08 | 41.90 | 65.25 |
|  | 0.3 | **89.62** | 42.78 | **69.00** |
|  | 0.5 | 89.24 | 41.97 | 66.43 |
|  | 0.7 | 89.46 | **43.59** | 67.27 |
|  | 1 | 89.30 | 42.12 | 66.82 |

Note: $\alpha$ and $\gamma$ are the coefficients of focal loss, and $\beta$ is the coefficient of auxiliary task. Bold indicates the best result for each coefficient.

As shown in the fourth row of Table 3, the introduction of the auxiliary task significantly improved the precision of the model by 9.49%, recall by 11.49%, accuracy by 1.85%. This indicates that the introduction of the auxiliary task of regression prediction of activity can help the model better learn the features of compound-protein interaction through the way of influencing gradient descent. We also tested different β values in Table 4 and found that good results were obtained at 0.7.

### 4.2 Decoding strategy study

In order to better simulate compound-protein interactions in the decoder, we attempted different decoding methods in the decoder

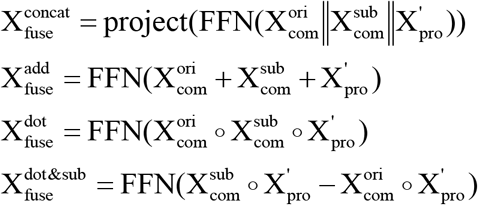

where ○ represents the Hadamard product. We found that the concatenation method achieved better overall performance with an accuracy of 89.33%, recall of 43.15% and precision of 66.52%.

### 4.3 Pretrained embedding effect study

In F-CPI (M), we use pre-trained embedding with FFN to extract protein sequence features, while for compound sequence features, we use learnable word embedding with self-attention encoder layers for feature extraction. In Table 6, we tested the model’s performance when pre-trained embedding was used for compounds and when pre-trained word embeddings were not used for proteins. The pre-trained embedding features of compounds are derived from chemBERTa. We also calculated the training time under the same hyperparameters and found that although introducing pre-trained compound embeddings only took half of the time, it significantly reduced the accuracy of the model by 0.93%, recall by 6.62%, precision by 3.97%. On the other hand, introducing protein pre-trained embeddings in F-CPI(M) not only significantly reduced the training time by about 80% but also achieved better accuracy by 0.09%, recall 0.22%, precision by 0.57% compared to the learnable word embedding layer.

**Table 5.** Performance of using different decode method.

| Methods | ACC(%) | RECALL(%) | PRE(%) |
| --- | --- | --- | --- |
| dot&sub | 89.26 | 38.14 | **68.88** |
| dot | 88.37 | 30.49 | 65.40 |
| add | 87.80 | 35.13 | 58.46 |
| concat | **89.46** | **43.59** | 67.27 |

Note: sub represents subtraction operation, concat represents concatenation operation.
Bold font indicates the best result.

**Table 6.** Study of pretrained embedding.

| Methods | ACC(%) | RECALL(%) | PRE(%) | TIME(min) |
| --- | --- | --- | --- | --- |
| w p-emb(both) | 88.53 | 36.97 | 63.30 | **258** |
| w/o p-emb(both) | 89.37 | 43.37 | 66.70 | 2317 |
| base | **89.46** | **43.59** | **67.27** | 489 |

Note: both means that both the compound and the protein used (or not used) pre-trained word embedding methods. Bold font indicates the best result.

For this phenomenon, we attribute it to the complexity of protein structures. Pre-trained embeddings can better extract their hidden high-dimensional features from sequence features. However, for compounds, due to the high similarity between original compounds and F-substituted compounds, pre-trained embedding methods trained on large datasets cannot distinguish the small disturbances caused by F-substitution well. In the pre-trained embedding layer, they focus too much on similarity and lose some information, which is crucial for CPI-FS tasks. This is also one of the reasons why directly using the regression CPI model described above cannot achieve satisfactory results.

## 5. Case study

In the previous text, we validated the superiority of the F-CPI model in F-substitution problems on the constructed dataset. In order to further explore the potential of the model in practical applications, we designed a series of wet experiments to verify the model’s ability to assist in drug optimization decision-making.

### 5.1 Deep learning models and algorithm applications

For a specific drug molecule, first, all hydrogen atoms of the molecule are sequentially replaced with fluorine atoms to obtain n candidate molecules, which are then combined with the original molecule and target protein to obtain n candidate triplets (F_ori_, F_sub_, P) that match the input of the previous model, where n is the number of hydrogen atoms in the original molecule. Then, model try to predict the candidate triplets. Among the m positive samples provided by the model, we ranked them based on the probability of the final binary vector output by the model, and selected the top 10 candidate molecules. Finally, we measured the difficulty of molecular retrosynthesis and selected specific molecules for synthesis and wet experiments to verify the model’s judgment.

It is worth mentioning that the selected model is the w/o pssm version, which takes into account the universality of the model, that is, some unknown proteins that have not appeared during training may fail in multiple sequence alignment. At the same time, as shown in Table 3, the w/o pssm model has almost the same performance as the original model. In order to further improve the model’s ability in application, we merged the test set from the previous testing process into the training set, retaining only the validation set as the basis for selecting epochs for training. Finally, we selected appropriate model parameters based on the highest recall criterion, in order to obtain more choices and enable us to conduct the final experiment while considering the difficulty and cost of reverse synthesis. We selected three pairs of compounds for the experiment.

### 5.2 Assay protocol

A fluorescence-based enzyme inhibition assay in 96-well plate format was used to assess the inhibition activity of SARS-CoV-2 3CLpro (6×His). The hydrolytic rates of Dabcyl-KNSTLQSGLRKE-Edans (DKE) were monitored in a 100 μL reaction mixture. Briefly, the SARS-CoV-2 3CLpro (6×His) was preincubated with analytes in 90 μL reaction buffer at 37 °C for 60 min. The reaction buffer included 1 × PBS, 1 mM EDTA. Then the hydrolytic reaction was proceeded for 20 min by the addition of 10 μL DKE. The final concentration of enzyme and DKE were 4 μg/mL and 20 μM, respectively. The generated fluorescent signals (excitation/emission, 340 nm/490 nm) were monitored by the microplate reader (SpectraMax® iD3, Molecular Devices, Austria)[43-45].

### 5.3 Positive sample cases

As show in Figure 5, in case (a), we perform an F-substitution operation on (1*R*,2*S*,5*S*)-3-((S)-2-acetamido-2-(3,5-dichloro-4-(2-(dimethylamino)ethoxy)phenyl)acetyl)-*N*-((*S*)-1-cyano-2-((*S*)-2-oxopyrrolidin-3-yl)ethyl)-6,6-dimethyl-3-azabicyclo[3. 1.0]hexane-2-carboxamide (**1**) to obtain (1*R*,2*S*,5*S*)-*N*-((*S*)-1-cyano-2-((*S*)-2-oxopyrrolidin-3-yl)ethyl)-3-((*S*)-2-(3,5-dichloro-4 -(2-(dimethylamino)ethoxy)phenyl)-2-(2-fluoroacetamido)acetyl)-6,6-dimethyl-3-aza bicyclo[3.1.0]hexane-2-carboxamide (**2**) (Figure S1-S2). After F-substitution, The IC50 value decreased from 28.19 to 0.23. This significant increase in activity was consistent with the model’s prediction, which effectively verified the predictive ability of the model (Table S1).

**Figure 5.**
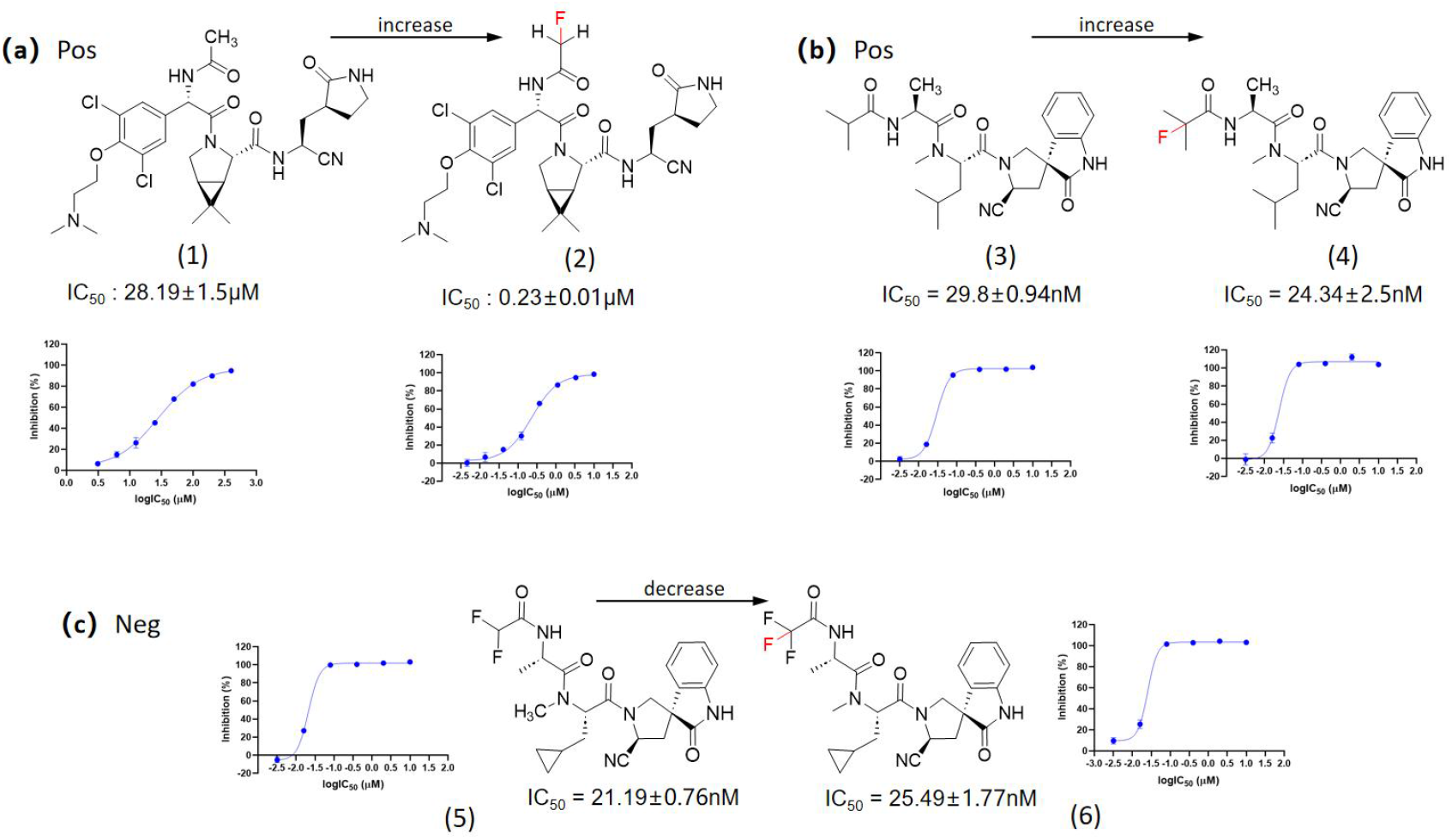
positive cases (a) (b) and negative case (c). The top of the case shows the structural formula before and after F-substitution, and the bottom shows the results measured in experiments.

In case (b), we perform an F-substitution operation on (*S*)-*N*-((*S*)-1-((3*R*,5’*S*)-5’-cyano-2-oxospiro[indoline-3,3’-pyrrolidin]-1’-yl)-4-methyl-1 -oxopentan-2-yl)-2-isobutyramido-*N*-methylpropanamide (**3**) to obtain (*S*)-*N*-((*S*)-1-((3*R*,5’*S*)-5’-cyano-2-oxospiro[indoline-3,3’-pyrrolidin]-1’-yl)-4-methyl-1 -oxopentan-2-yl)-2-(2-fluoro-2-methylpropanamido)-*N*-methylpropanamide (**4**) (Figure S3-S4).

After F-substitution, The IC_50_ value decreased from 29.80 to 24.34, indicating an increase in drug activity, which is consistent with the model prediction (Table S1).

### 5.4 Negative sample cases

As show in Figure 5, in case (c), based on the previous selection principle, we selected a pair of negative samples as a negative control, we perform an F-substitution operation on (*S*)-*N*-((*S*)-1-((3*R*,5’*S*)-5’-cyano-2-oxospiro[indoline-3,3’-pyrrolidin]-1’-yl)-3-cyclopropyl-1-oxopropan-2-yl)-2-(2,2-difluoroacetamido)-*N*-methylpropanamide (**5**) to obtain (*S*)-*N*-((*S*)-1-((3*R*,5’*S*)-5’-cyano-2-oxospiro[indoline-3,3’-pyrrolidin]-1’-yl)-3-cyclopropyl-1-oxopropan-2-yl)-*N*-methyl-2-(2,2,2-trifluoroacetamido)propenamide (**6**) (Figure S5-S6).

After F-substitution, The IC_50_ value increased from 21.19 nM to 25.49 nM, indicating a decrease in drug activity, which is consistent with the model prediction (Table S2).

## Conclusion

In order to investigate the effect of F-substitution on activity changes, we compiled the largest dataset currently containing 111,168 samples, designed a specialized model pattern F-CPI, and compared multiple methods horizontally. We found that the F-CPI model achieved satisfactory performance overall. In the end, we applied F-CPI to the structural optimization of hit compounds against SARS-CoV-2 3CL_pro_. Impressively, in one case, the introduction of only one F atom, the activity increased by more than two orders of magnitude (IC_50_: 22.99 nM vs. 28190 nM). F-CPI has certain practical significance and is expected to provide new ideas for the discovery of fluorinated drugs.

## Supporting Information

Table S1: IC_50_ and SE of drug molecules in positive case; Table S2: IC_50_ and SE of drug molecules in negative case; Figure S1: The synthesis path of (**1**); Figure S2: The synthesis path of (**2**); Figure S3: The synthesis path of (**3**); Figure S4: The synthesis path of (**4**); Figure S5: The synthesis path of (**5**); Figure S6: The synthesis path of (**6**).

## Acknowledgments

The authors sincerely acknowledge the financial support from the National Natural Science Foundation of China (82322067), the National Key Research and Development Program of China (2022YFA1004304) and the funds from Shanghai Institute of Materia Medica (SIMM0120231003).

## Data and Software Availability

the core dataset and python scripts are available at the following github repository: https://github.com/ywwhhh/F-CPI.

