## Supplementary material for "F-CPI: Prediction of activity changes induced by fluorine substitution using multimodal deep learning": Table S1; Table S2; Figure S1; Figure S2; Figure S3; Figure S4; Figure S5; Figure S6.

**Table S1** $IC_{50}$ and SE of drug molecules in positive case

| Comp. | IC50 (μM) | SE (μM) | incubation time (min) |
|---|---|---|---|
| 1 | 28.19 | 1.50 | 60 |
| **2** | **0.23** | **0.01** | **60** |
| 3 | 29.80 | 0.94 | 60 |
| **4** | **24.34** | **2.50** | **60** |

Note: Bold font indicates the F-substituted compound.

**Table S2** $IC_{50}$ and SE of drug molecules in negative case

| Comp. | $IC_{50}$ (nM) | SE (nM) | incubation time (min) |
|---|---|---|---|
| 5 | 21.19 | 0.76 | 60 |
| **6** | **25.49** | **1.77** | **60** |

Note: Bold font indicates the original compound.

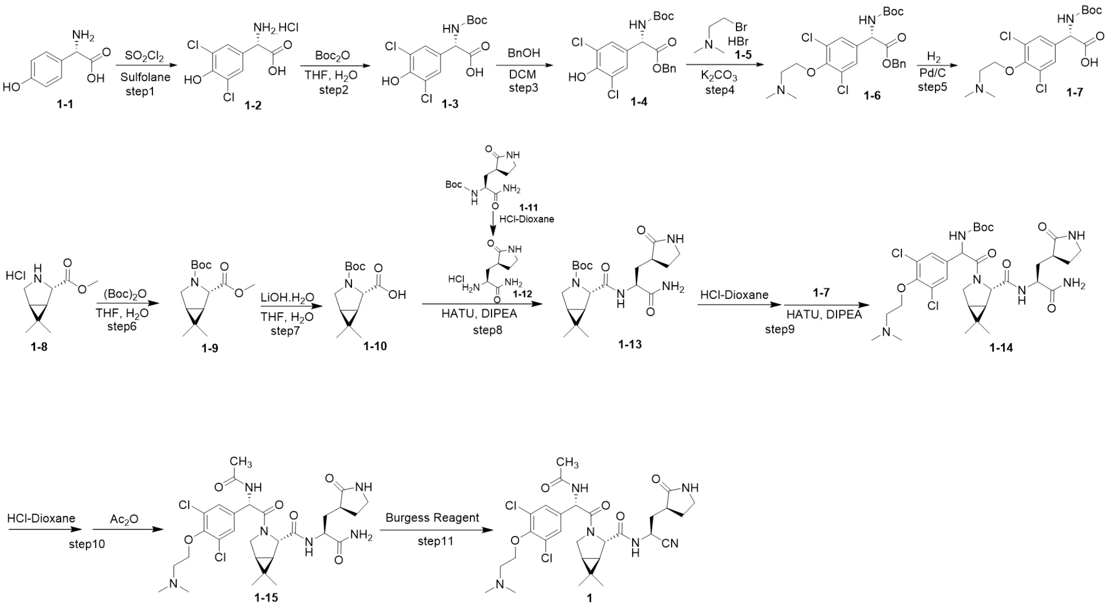


**Figure S1** The synthesis route of **1**.

**(*S*)-2-amino-2-(3,5-dichloro-4-hydroxyphenyl)acetic acid hydrochloride (1-2).** Compound **1-1** (25 g, 149.6 mmol) was dissolved in sulfolane (250 mL), sulfonyl chloride (30 mL, 373.9 mmol) was added dropwise to the reaction at 40 °C and the reaction was stirred at 60 °C for 2 h. Then toluene (400 mL) added to the reaction, the mixture was filtered to give compound **1-2** as white solid (34 g, yield: 83.5%). $^{1}H$ NMR (400 MHz, DMSO-*d*$_6$) δ 10.61 (s, 1H), 8.93 (s, 3H), 7.56 (s, 2H), 5.08 (s, 1H). ESI-MS m/z 236.02 $[M+H]^+$.

**(*S*)-2-((*tert*-butoxycarbonyl)amino)-2-(3,5-dichloro-4-hydroxyphenyl)acetic acid (1-3).** Compound **1-2** (34 g, 124.8 mmol) was dissolved in THF (350 mL) and $H_2O$ (350 mL), sodium carbonate (66 g, 623.8 mmol) and $(Boc)_2O$ (43 mL, 187.2 mmol) were added to the reaction at 0 °C, the reaction was stirred at 25 °C for 5 h. Then EA (50 mL) and 1N HCl aqueous solution (10 mL) were added to the reaction. After separation, the organic phase was washed with saturated brine (10 mL), dried over anhydrous sodium sulphate, concentrated under vacuum and purified by column chromatography to give compound **1-3** as white solid (32 g, yield: 76.2%). $^{1}H$ NMR (400 MHz, $CDCl_3$) δ 8.02 (d, J = 5.0 Hz, 1H), 7.36 (s, 2H), 7.30 (d, J = 13.5 Hz, 1H), 5.01 (d, J = 5.0 Hz, 1H), 1.27 (d, J = 11.6 Hz, 9H).

**Benzyl (*S*)-2-((*tert*-butoxycarbonyl)amino)-2-(3,5-dichloro-4-hydroxyphenyl) acetate (1-4).** Compound **1-3** (30 g, 89.2 mmol) was dissolved in DCM (300 mL), benzyl alcohol (14 mL, 133.9 mmol), EDCI (21.4 g, 111.5 mmol) and DMAP (16.3 g, 133.9 mmol) were added to the reaction at 0 °C, the reaction was stirred at 25 °C for 10 h. Then DCM (50 mL) and 1*N* HCl aqueous solution (10 mL) were added to the reaction. After separation, the organic phase was washed with water (10 mL) and saturated brine (10 mL), dried over anhydrous sodium sulphate, concentrated under vacuum and purified by column chromatography to give compound **1-4** as white solid (13.9 g, yield: 36.7%).$^{1}$H NMR (400 MHz, DMSO-$d_6$) δ 10.24 (s, 1H), 7.84 (d, J = 8.1 Hz, 1H), 7.42 (s, 2H), 7.31 (q, J = 4.1, 3.1 Hz, 3H), 7.28-7.21 (m, 2H), 5.24 (d, J = 8.0 Hz, 1H), 5.18-5.08 (m, 2H), 1.38 (s, 9H).

**Benzyl (*S*)-2-((*tert*-butoxycarbonyl)amino)-2-(3,5-dichloro-4-(2-(dimethylamino)ethoxy) phenyl) acetate (1-6).** Compound **1-4** (13 g, 30.6 mmol) was dissolved in acetone (130 mL), potassium carbonate (12.6 g, 91.8 mmol) and compound **1-5** (10.6 g, 45.9 mmol) were added to the reaction, the reaction was stirred at 40 °C for 10 h. Filter the reaction solution, EA (50 mL) and water (10 mL) were added to the reaction. After separation, the organic phase was washed saturated brine (10 mL), dried over anhydrous sodium sulphate, concentrated under vacuum and purified by column chromatography to give compound **1-6** as yellow oil (3.6 g, yield: 15.2%). $^{1}$H NMR (400 MHz, $CDCl_3$) δ 7.31 (dd, $J$ = 4.8, 1.9 Hz, 3H), 7.26 (s, 2H), 7.20 (dd, $J$ = 6.7, 2.9 Hz, 2H), 5.65 (d, $J$ = 7.1 Hz, 1H), 5.25 (d, $J$ = 7.1 Hz, 1H), 5.15 (s, 2H), 4.13 (t, $J$ = 5.6 Hz, 2H), 2.90 (t, $J$ = 5.6 Hz, 2H), 2.46 (s, 6H), 1.41 (s, 9H).

**(*S*)-2-((*tert*-butoxycarbonyl)amino)-2-(3,5-dichloro-4-(2-(dimethylamino)ethoxy)phenyl) acetic acid (1-7).** Compound **1-6** (3.4 g, 8.2 mmol) was dissolved in MeOH (34 mL), 5% Pd/C (340 mg) were added to the reaction, the reaction was stirred under $H_2$ atmosphere at 25 °C for 2 h. Filtration of reaction solution, concentrated under vacuum to give compound **1-7** as white solid (2.7 g, yield: 93.1%). ESI-MS m/z 407.07 $[M+H]^+$.

**(1*R*,2*S*,5*S*)-3-(*tert*-butoxycarbonyl)-6,6-dimethyl-3-azabicyclo[3.1.0]hexane-2-carboxylic acid (1-10).** Compound **1-8** (6.0 g, 29.2 mmol) was dissolved in THF (60 mL) and $H_2O$ (60 mL), sodium hydroxide (2.3 g, 58.4 mmol) and $(Boc)_2O$ (8 mL, 35.0 mmol) were added to the reaction at 0 °C, the reaction was stirred at 25 °C for 2 h. Then EA (20 mL) and 1*N* HCl aqueous solution (5 mL) were added to the reaction. After separation, the organic phase was washed with saturated sodium bicarbonate (5 mL), and saturated brine (5 mL), dried over anhydrous sodium sulphate, concentrated under vacuum and purified by column chromatography to give 6.0 g crude compound **1-9** as yellow oil. THF (60 mL) and $H_2O$ (60 mL) were added to compound **1-9** solution, then lithium hydroxide monohydrate (3.7 g, 89.1 mmol) was added to the reaction, the reaction was stirred at 40 °C for 2 h. Then EA (30 mL) and 1*N* HCl aqeous solution (10 mL) were added to the reaction. After separation, the organic phase was washed with saturated sodium bicarbonate (5 mL), and saturated brine (5 mL), dried over anhydrous sodium sulphate, concentrated under vacuum and purified by column chromatography to give 3.3 g compound **1-10** as yellow oil. $^1H$ NMR (400 MHz, DMSO-$d_6$) δ 12.62 (s, 1H), 3.90 (d, J = 15.5 Hz, 1H), 3.52-3.43 (m, 1H), 3.27 (dd, J = 13.9, 10.9 Hz, 1H), 1.41-1.38 (m, 2H), 1.34 (d, J = 19.3 Hz, 9H), 1.00 (s, 3H), 0.90 (d, J = 6.1 Hz, 3H).

***tert*-Butyl (1*R*,2*S*,5*S*)-2-(((*S*)-1-amino-1-oxo-3-((*S*)-2-oxopyrrolidin-3-yl)propan-2-yl)carbamoyl)-6,6-dimethyl-3-azabicyclo[3.1.0]hexane-3-carboxylate (1-13).** Compound **1-11** (3.2 g, 12 mmol) and 4 M HCl in 1,4-dioxane (30 mL) were dissolved in 1,4-dioxane (30 mL), the reaction was stirred at 40 °C for 1 h. And then the solution was concentrated to remove the solvent, dry HCl salt **1-12** was obtained. On the other hand, compound **1-10** (3.0 g, 12.0 mmol) and HATU (6.8 g, 18 mmol) were dissolved in DCM (35 mL), the reaction was stirred at 25 °C for 0.5 h. Then DIPEA (4.6 mL, 26.4 mmol) and **1-12** were added to the reaction, the reaction mixture was stirred for 10 h at 25 °C. Then DCM (20 mL) and 1N HCl aqueous solution (5 mL) were added to the reaction. After separation, the organic phase was washed with water (5 mL) and saturated brine (5 mL), dried over anhydrous sodium sulphate, concentrated under vacuum and purified by column chromatography to give compound **1-13** as white solid (4.0 g, yield: 82.9%). $^1H$ NMR (400 MHz, DMSO-$d_6$) δ 8.18 (dd, J = 11.4, 8.3 Hz, 1H), 7.61 (d, J = 16.0 Hz, 1H), 7.39-7.19 (m, 1H), 7.07-6.99 (m, 1H), 4.26 (ddt, J = 12.6, 8.5, 4.3 Hz, 1H), 4.01 (d, J = 18.0 Hz, 1H), 3.54 (ddd, J = 35.6, 10.8, 4.3 Hz, 1H), 3.27 (dd, J = 10.9, 6.8 Hz, 1H), 3.23-3.00 (m, 2H), 2.39-2.10 (m, 2H), 1.96 (dddd, J = 15.7, 13.6, 11.4, 3.9 Hz, 1H), 1.73 (dq, J = 12.4, 9.1 Hz, 1H), 1.52 (dddd, J = 21.5,

18.1, 9.8, 5.6 Hz, 1H), 1.37-1.28 (m, 9H), 1.28-1.23 (m, 2H), 1.00 (s, 3H), 0.89 (s, 3H).

***tert*-Butyl (2-((1*R*,2*S*,5*S*)-2-(((*S*)-1-amino-1-oxo-3-((*S*)-2-oxopyrrolidin-3-yl)propan-2-yl) carbamoyl)-6,6-dimethyl-3-azabicyclo[3.1.0]hexan-3-yl)-1-(3,5-dichloro-4-(2-(dimethylamino) ethoxy)phenyl)-2-oxoethyl)carbamate (1-14).** Compound 1-13 (2.6 g, 6.4 mmol) and 4 *N* HCl in 1,4-dioxane (16 mL) were dissolved in 1,4-dioxane (16 mL), the reaction was stirred at 40 °C for 1 h. And then the solution was concentrated to remove the solvent, dry HCl salt was obtained. On the other hand, compound 1-7 (2.6 g, 12.0 mmol) and HATU (3.6 g, 9.6 mmol) were dissolved in DCM (35 mL), the reaction was stirred at 25 °C for 0.5 h. Then DIPEA (2.5 mL, 14.1 mmol) and HCl salt were added to the reaction, the reaction mixture was stirred for 10 h at 25 °C. Then DCM (6 mL) and sodium hydrogen carbonate (2 mL) were added to the reaction, dried over anhydrous sodium sulphate, concentrated under vacuum and purified by column chromatography to give compound 1-14 as yellow solid (3.2 g, yield: 52.3%). ESI-MS m/z 697.37 $[M+H]^+$.

**(1*R*,2*S*,5*S*)-3-((S)-2-acetamido-2-(3,5-dichloro-4-(2-(dimethylamino)ethoxy)phenyl)acetyl)-*N*-((*S*)-1-cyano-2-((*S*)-2-oxopyrrolidin-3-yl)ethyl)-6,6-dimethyl-3-azabicyclo[3.1.0]hexane-2-carboxamide (1).** Compound 1-14 (400 mg, 0.57 mmol) and 4 *N* HCl in 1,4-dioxane (1 mL) were dissolved in 1,4-dioxane (3 mL), the reaction was stirred at 40 °C for 1 h. Then the reaction was concentrated to remove the solvent and $Et_3N$ (0.2 mL, 2.7 mmol), acetic anhydride (0.2 mg, 1.12 mmol), DCM (3 mL) were added to the reaction at 0 °C. The reaction was allowed to warm to 25 °C and stirred for 10 h. Then DCM (6 mL) and sodium hydrogen carbonate (2 mL) were added to the reaction, dried over anhydrous sodium sulphate, concentrated under vacuum and purified by column chromatography to give 200 mg crude compound 1-15 as yellow solid. Compound 1-15 and Burgess Reagent (166 mg, 0.7 mmol) were dissolved in dried DCM (3 mL), the reaction was stirred at 25 °C for 10 h. The mixture was concentrated and purified by column chromatography to give 15 mg compound 1 as white solid. $^1H$ NMR (600 MHz, $CDCl_3$) δ 8.75 (d, J = 5.7 Hz, 1H), 7.87 (s, 2H), 7.60 (s, 2H), 6.53 (s, 1H), 4.55 (dt, J = 11.0, 5.3 Hz, 1H), 4.47 (td, J = 5.8, 3.5 Hz, 2H), 4.17 (s, 1H), 4.07 (dd, J = 9.5, 4.0 Hz, 1H), 3.47 (s, 1H), 3.40 (td, J = 10.1, 9.6, 7.3 Hz, 2H), 3.21 (d, J = 7.3 Hz, 2H), 3.16 (s, 6H), 3.02 (d, J = 9.6 Hz, 1H), 2.38 (s, 3H), 2.19 (tq, J = 11.3, 8.2

Hz, 2H), 2.02 (s, 2H), 1.98-1.90 (m, 1H), 1.82 (dq, J = 12.7, 9.5 Hz, 1H), 1.08 (d, J = 2.8 Hz, 6H).

ESI-MS m/z 621.18 $[M+H]^+$.

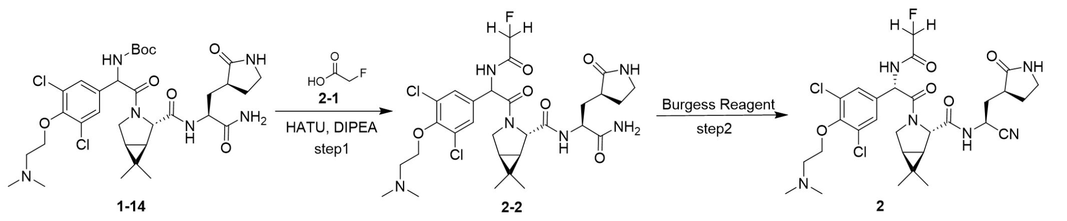


**Figure S2** The synthesis route of **2**.

**(1*R*,2*S*,5*S*)-*N*-((*S*)-1-cyano-2-((*S*)-2-oxopyrrolidin-3-yl)ethyl)-3-((*S*)-2-(3,5-dichloro-4-(2-(dimethylamino)ethoxy)phenyl)-2-(2-fluoroacetamido)acetyl)-6,6-dimethyl-3-azabicyclo[3.1.0]hexane-2-carboxamide (2).** Compound 1-14 (400 mg, 0.57 mmol) and 4 *N* HCl in 1,4-dioxane (1.2 mL) were dissolved in 1,4-dioxane (1.2 mL), the reaction was stirred at 40 °C for 1 h. And then the solution was concentrated to remove the solvent, dry HCl salt was obtained. On the other hand, compound 2-1 (44 mg, 0.57 mmol) and HATU (325 mg, 0.86 mmol) were dissolved in DCM (5 mL) under $N_2$ atmosphere, the reaction was stirred at 25 °C for 0.5 h. Then DIPEA (0.2 mL, 1.25 mmol), and HCl salt were added to the reaction, the reaction mixture was stirred for 10 h at 25 °C. Then DCM (6 mL) and sodium hydrogen carbonate (2 mL) were added to the reaction, dried over anhydrous sodium sulphate, concentrated under vacuum and purified by column chromatography to give 200 mg crude compound 2-2 as yellow solid. 2-2 and Burgess Reagent (285 mg, 1.2 mmol) were dissolved in dried DCM (3 mL), the reaction was stirred at 25 °C for 10 h. The mixture was concentrated and purified by column chromatography to give 20 mg compound 2 as white solid. $^{1}$H NMR (400 MHz, DMSO-$d_6$) δ 9.04 (d, J = 8.0 Hz, 1H), 8.79-8.64 (m, 1H), 7.69 (s, 1H), 7.62-7.46 (m, 2H), 5.68 (d, J = 7.1 Hz, 1H), 4.91 (d, J = 9.9 Hz, 1H), 4.79 (d, J = 9.8 Hz, 1H), 4.41 (s, 1H), 4.21 (d, J = 20.2 Hz, 2H), 3.89 (d, J = 6.5 Hz, 1H), 3.51 (d, J = 33.6 Hz, 2H), 3.17-3.00 (m, 1H), 2.78 (s, 3H), 2.56-2.44 (m, 6H), 2.13 (s, 2H), 1.71 (dd, J = 20.1, 10.3 Hz, 1H), 1.54 (dd, J = 17.2, 6.2 Hz, 2H), 1.30 (d, J = 7.5 Hz, 1H), 1.25- 1.15 (m, 1H), 0.99 (d, J = 32.3 Hz, 6H). ESI-MS m/z 639.48 [M+H].

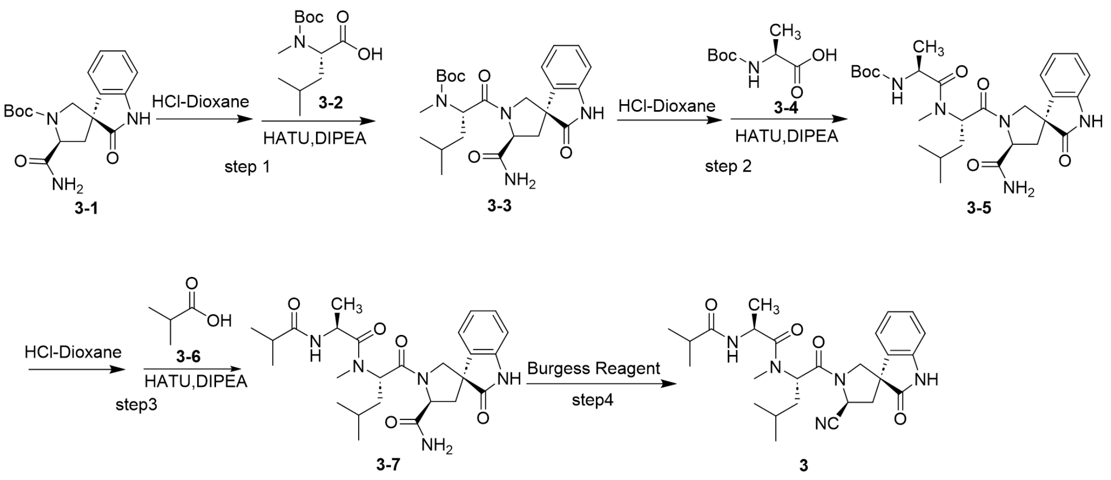


**Figure S3** The synthesis route of 3.

***tert*-butyl((*S*)-1-((3*R*,5'*S*)-5'-carbamoyl-2-oxospiro[indoline-3,3'-pyrrolidin]-1'-yl)-4-methyl-1-oxopentan-2-yl)(methyl)carbamate (3-3).** Compound 3-1 (800 mg, 2.42 mmol) and 4 *M* HCl in 1,4-dioxane (6 mL) were dissolved in 1,4-dioxane (6 mL), the reaction was stirred at 40 °C for 1 h. And then the solution was concentrated to remove the solvent to obtain white solid. Compound 3-2 (600 mg, 2.44 mmol) and HATU (1376 mg, 3.62 mmol) were dissolved in DCM (20 mL) under $N_2$ atmosphere, the reaction was stirred at 25 °C for 0.5 h. Then DIPEA (1.0 mL, 5.64 mmol), and white solid were added to the reaction, the reaction mixture was stirred for 10 h at 25 °C. Then DCM (10 mL) and 1*N* HCl aqueous solution (2 mL) were added to the reaction. After separation, the organic phase was washed with water (2 mL) and saturated brine (2 mL), dried over anhydrous sodium sulphate, concentrated under vacuum and purified by column chromatography to give compound 3-3 as off-white solid (900 mg, yield: 81.3%). $^{1}$H NMR (400 MHz, DMSO-$d_6$) δ 10.69 (d, J = 12.1 Hz, 1H), 7.50 (s, 1H), 7.23 (t, J = 7.7 Hz, 1H), 7.06 (s, 1H), 6.97 (t, J = 7.5 Hz, 1H), 6.92-6.74 (m, 2H), 4.87-4.49 (m, 2H), 3.97-3.63 (m, 2H), 2.77-2.60 (m, 3H), 2.37-2.13 (m, 2H), 1.50-1.30 (m, 2H), 1.30-1.20 (m, 1H), 1.05 (d, J = 3.0 Hz, 9H), 0.87 (dd, J = 11.9, 6.5 Hz, 6H). ESI-MS m/z 458.56 $[M+H]^+$.

***tert*-butyl((*S*)-1-(((*S*)-1-((3*R*,5'*S*)-5'-carbamoyl-2-oxospiro[indoline-3,3'-pyrrolidin]-1'-yl)-4-methyl-1-oxopentan-2-yl)(methyl)amino)-1-oxopropan-2-yl)carbamate (3-5).** Compound **3-3** (430 mg, 0.93 mmol) and 4 *M* HCl in 1,4-dioxane (3 mL) were dissolved in 1,4-dioxane (3 mL), the reaction was stirred at 40 °C for 1 h. And then the solution was concentrated to remove the solvent, white solid was obtained. On the other hand, compound **3-4** (178 mg, 0.94 mmol) and HATU (535 mg, 1.41 mmol) were dissolved in DCM (10 mL) under $N_2$ atmosphere, the reaction was stirred at 25 °C for 0.5 h. Then DIPEA (0.4 mL, 2.1 mmol), and white solid were added to the reaction, the reaction mixture was stirred for 10 h at 25 °C. Then DCM (5 mL) and 1N HCl aqueous solution (1 mL) were added to the reaction. After separation, the organic phase was washed with water (2 mL) and saturated brine (2 mL), dried over anhydrous sodium sulphate, concentrated under vacuum and purified by column chromatography to give **3-5** as white solid (310 mg, yield: 62.5%).$^1$H NMR (400 MHz, DMSO-$d_6$) δ 10.72 (s, 1H), 7.53 (s, 1H), 7.28-7.19 (m, 1H), 7.06 (s, 1H), 6.99 (t, J = 7.5 Hz, 1H), 6.92- 6.79 (m, 3H), 5.22 (dd, J = 9.8, 5.2 Hz, 1H), 4.64- 4.51 (m, 1H), 4.17- 4.00 (m, 1H), 3.77- 3.54 (m, 2H), 2.93- 2.78 (m, 3H), 2.36- 2.05 (m, 2H), 1.42-1.19 (m, 15H), 0.96- 0.73 (m, 6H). ESI-MS m/z 529.85 $[M+H]^+$

***(3R,5'S)-1'-(N-(isobutyryl-L-alanyl)-N-methyl-L-leucyl)-2-oxospiro[indoline-3,3'-pyrrolidine]-5'-carboxamide (3-7).*** Compound **3-5** (250 mg, 0.47 mmol) and 4 *N* HCl in 1,4-dioxane (3 mL) were dissolved in 1,4-dioxane (3 mL), the reaction was stirred at 40 °C for 1 h. And then the solution was concentrated to remove the solvent, dry HCl salt was obtained. On the other hand, compound **3-6** (42 mg, 0.47 mmol) and HATU (270 mg, 0.71 mmol) were dissolved in DCM (4 mL) under $N_2$ atmosphere, the reaction was stirred at 25 °C for 0.5 h. Then DIPEA (0.2 mL, 1.04 mmol), and HCl salt were added to the reaction, the reaction mixture was stirred for 10 h at 25 °C. Then DCM (5 mL) and 1*N* HCl aqueous solution (0.5 mL) were added to the reaction. After separation, the organic phase was washed with water (1 mL) and saturated brine (1 mL), dried over anhydrous sodium sulphate, concentrated under vacuum and purified by column chromatography to give compound **3-7** as white solid (195 mg, yield: 83.0%). $^1$H NMR (400 MHz, DMSO-$d_6$) δ 10.72 (s, 1H), 7.85 (d, *J* = 8.0 Hz, 1H), 7.52 (s, 1H), 7.23 (t, *J* = 7.7 Hz, 1H), 7.06 (s, 1H), 6.99 (t, *J* = 7.6 Hz, 1H), 6.90 (d, *J* = 7.7 Hz, 1H), 6.81 (d, *J* = 7.4 Hz, 1H), 5.21 (dd, *J* = 9.3, 5.4 Hz, 1H), 4.56 (t, *J* = 9.0 Hz, 1H), 4.35 (p, *J* = 6.9 Hz, 1H), 3.74-3.63 (m, 2H), 2.84 (s, 3H), 2.29 (ddd, *J* = 29.1, 13.3, 7.6 Hz, 2H), 2.15 (dd, *J* = 12.5, 10.0 Hz, 1H), 1.55 (t, *J* = 10.0 Hz, 1H), 1.48-1.31 (m, 1H), 1.28 (dd, *J* = 11.9, 6.5 Hz, 4H), 0.92-0.84 (m, 6H), 0.80 (d, *J* = 6.0 Hz, 3H), 0.42 (d, *J* = 6.8 Hz, 3H). ESI-MS m/z 500.3 $[M+H]^+$.

**(*S*)-*N*-((*S*)-1-((3*R*,5'*S*)-5'-cyano-2-oxospiro[indoline-3,3'-pyrrolidin]-1'-yl)-4-methyl-1-oxopentan-2-yl)-2-isobutyramido-*N*-methylpropanamide (3).** Compound **3-7** (100 mg, 0.2 mmol) was dissolved in dried DCM (4 mL) under $N_2$ atmosphere. Burgess Reagent (95 mg, 0.4 mmol) was added and stirred at 25 °C for 10 h. The mixture was concentrated and purified by column chromatography to give the compound **3** as white solid (30 mg, yield: 31%). $^1$H NMR (400 MHz, $CDCl_3$) δ 8.77 (s, 1H), 7.26 (td, *J* = 7.8, 1.2 Hz, 1H), 7.04- 6.98 (m, 1H), 6.94 (d, *J* = 7.8 Hz, 1H), 6.80 (dd, *J* = 7.5, 1.1 Hz, 1H), 5.33- 5.27 (m, 1H), 4.97 (dd, *J* = 9.2, 8.2 Hz, 1H), 4.75 (dd, *J* = 7.6, 6.7 Hz, 1H), 4.12 (q, *J* = 7.1 Hz, 1H), 4.02 (dd, *J* = 10.5, 1.3 Hz, 1H), 3.95 (d, *J* = 10.5 Hz, 1H), 3.09 (s, 3H), 2.84 (dd, *J* = 13.2, 9.3 Hz, 1H), 2.51 (ddd, *J* = 13.2, 8.2, 1.2 Hz, 1H), 2.29 (p, *J* = 6.9 Hz, 1H), 1.72 (td, *J* = 6.8, 6.0, 1.3 Hz, 2H), 1.08 (dd, *J* = 8.6, 6.9 Hz, 6H), 0.95 (d, *J* = 6.7 Hz, 3H), 0.89 (dd, *J* = 6.7, 5.7 Hz, 6H). $^{13}$C NMR (151 MHz, Chloroform-*d*) δ 176.35, 175.82, 173.74, 170.16, 139.97, 131.28, 129.67, 123.40, 122.24, 117.05, 110.89, 54.69, 52.92, 52.61, 46.16, 45.26, 39.19, 37.56, 35.50, 30.66, 24.89, 23.06, 22.28, 19.60, 19.42, 17.95. ESI-HRMS Calcd for $C_{26}H_{36}N_5O_4$ $[M+H]^+$: 482.2762, found 482.2751.

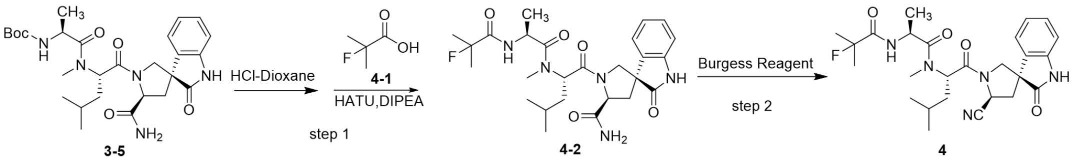

**Figure S4** The synthesis route of **4**.

**(3*R*,5'*S*)-1'-(*N*-((2-fluoro-2-methylpropanoyl)-*L*-alanyl)-*N*-methyl-*L*-leucyl)-2-oxospiro[indoline-3,3'-pyrrolidine]-5'-carboxamide (4-2).** Compound **3-5** (300 mg, 0.57 mmol) and 4 *N* HCl in 1,4-dioxane (3 mL) were dissolved in 1,4-dioxane (3 mL), the reaction was stirred at 40 °C for 1 h. And then the solution was concentrated to remove the solvent, white solid was obtained. On the other hand, compound **4-1** (60 mg, 0.57 mmol) and HATU (325 mg, 0.85 mmol) were dissolved in DCM (7 mL) under $N_2$ atmosphere, the reaction was stirred at 25 °C for 0.5 h. Then DIPEA (0.3 mL, 1.2 mmol), and HCl salt were added to the reaction, the reaction mixture was stirred for 10 h at 25 °C. Then DCM (5 mL) and 1*N* HCl aqueous solution (1 mL) were added to the reaction. After separation, the organic phase was washed with water (1 mL) and saturated brine (1 mL), dried over anhydrous sodium sulphate, concentrated under vacuum and purified by column chromatography to give compound **4-2** as white solid (190 mg, yield:64.8%). ESI-MS m/z 517.89 $[M+H]^+$

**(*S*)-*N*-((*S*)-1-((3R,5'S)-5'-cyano-2-oxospiro[indoline-3,3'-pyrrolidin]-1'-yl)-4-methyl-1-oxopentan-2-yl)-2-(2-fluoro-2-methylpropanamido)-*N*-methylpropanamide (4).** Compound **4-2** (170 mg, 0.33 mmol) was dissolved in dried DCM (4 mL) under $N_2$ atmosphere. Burgess Reagent (167 mg, 0.72 mmol) was added and stirred at 25 °C for 10 h. The mixture was concentrated and purified by column chromatography to give the compound **4** as white solid (70 mg, yield: 42.7%).$^1$H NMR (400 MHz, DMSO-$d_6$) δ 10.71 (s, 1H), 7.91 (dd, *J* = 7.4, 2.8 Hz, 1H), 7.23 (td, *J* = 7.5, 1.7 Hz, 1H), 7.07-6.93 (m, 2H), 6.88 (d, *J* = 7.7 Hz, 1H), 5.21 (dd, *J* = 9.5, 5.3 Hz, 1H), 5.14 (t, *J* = 8.0 Hz, 1H), 4.44 (td, *J* = 8.4, 7.8, 5.3 Hz, 1H), 3.76 (d, *J* = 10.6 Hz, 1H), 3.65 (d, *J* = 10.8 Hz, 1H), 2.88 (s, 3H), 2.68-2.59 (m, 1H), 2.45 (d, *J* = 7.3 Hz, 1H), 1.64 (ddt, *J* = 16.8, 11.9, 6.0 Hz, 1H), 1.50 (dt, *J* = 14.1, 9.2 Hz, 1H), 1.38 (dd, *J* = 21.8, 15.3 Hz, 6H), 1.25 (d, *J* = 9.0 Hz, 1H), 0.87 (d, *J* = 6.4 Hz, 3H), 0.80 (d, *J* = 6.2 Hz, 3H), 0.73 (d, *J* = 6.9 Hz, 3H). $^{13}$C NMR (101 MHz, DMSO-$d_6$) δ 176.98, 172.72, 172.19, 169.13, 141.82, 131.37, 129.33, 123.11, 122.54,

118.74, 110.40, 96.32, 54.60, 52.58, 52.51, 46.57, 45.42, 38.32, 36.94, 30.27, 25.31, 25.07, 24.07, 23.66, 22.36, 16.32. ESI-HRMS Calcd for $C_{26}H_{33}FN_5O_4$ $[M-H]^-$: 498.2517, found 498.2518.

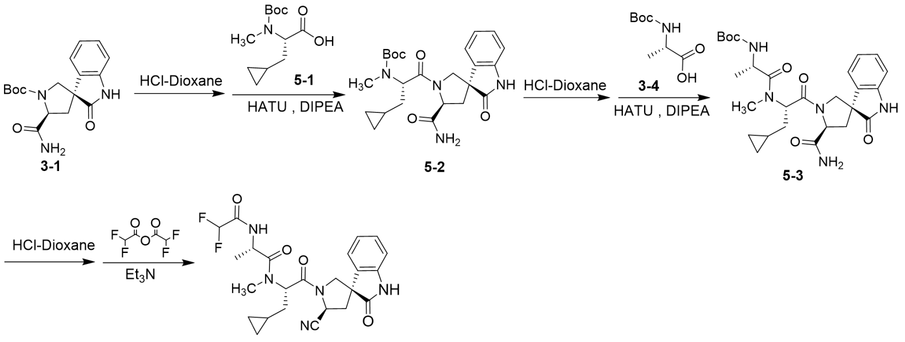


**Figure S5** The synthesis route of **5**.

***tert*-butyl((*S*)-1-((3*R*,5'*S*)-5'-carbamoyl-2-oxospiro[indoline-3,3'-pyrrolidin]-1'-yl)-3-cyclopropyl-1-oxopropan-2-yl)(methyl)carbamate (5-2).** Compound **3-1** (2.7 g, 8.2 mmol) and 4 M HCl in 1,4-dioxane (15 mL) were dissolved in 1,4-dioxane (15 mL), the reaction was stirred at 40 °C for 1 h. And then the solution was concentrated to remove the solvent, dry HCl salt was obtained. On the other hand, compound **5-1** (2.0 g, 8.2 mmol) and HATU (4.7 g, 12.3 mmol) were dissolved in DCM (40 mL) under $N_2$ atmosphere, the reaction was stirred at 25 °C for 0.5 h. Then DIPEA (3.0 mL, 18.0 mmol), and HCl salt were added to the reaction, the reaction mixture was stirred for 10 h at 25 °C. Then DCM (50 mL) and 1N HCl aqueous solution (10 mL) were added to the reaction. After separation, the organic phase was washed with water (10 mL) and saturated brine (10 mL), dried over anhydrous sodium sulphate, concentrated under vacuum and purified by column chromatography to give compound **5-2** as white solid (3.2 g, yield: 86.3%).$^1$H NMR (400 MHz, DMSO-$d_6$) δ 10.67 (s, 1H), 7.48 (s, 1H), 7.05 (d, J = 2.3 Hz, 1H), 6.97 (t, J = 7.6 Hz, 1H), 6.89 (d, J = 7.9 Hz, 2H), 6.82 (t, J = 7.8 Hz, 1H), 4.81 (dd, J = 9.0, 5.8 Hz, 1H), 4.54 (t, J = 8.8 Hz, 1H), 3.88 (t, J = 11.3 Hz, 1H), 3.70 (d, J = 10.3 Hz, 1H), 2.65 (s, 3H), 2.31-2.12 (m, 2H), 1.51-1.38 (m, 1H), 1.25-1.21 (m, 1H), 1.07 (s, 9H), 0.54 (t, J = 7.3 Hz, 1H), 0.33 (dd, J = 8.4, 5.0 Hz, 2H), 0.05 (td, J = 10.9, 8.3, 4.2 Hz, 2H). ESI-MS m/z 474.37 $[M+NH_4]^+$.

***tert*-butyl((*S*)-1-(((*S*)-1-((3*R*,5'*S*)-5'-carbamoyl-2-oxospiro[indoline-3,3'-pyrrolidin]-1'-yl)-3-cyclopropyl-1-oxopropan-2-yl)(methyl)amino)-1-oxopropan-2-yl)carbamate (5-3).** Compound **5-2** (2.3 g, 5.0 mmol) and 4 *N* HCl in 1,4-dioxane (15 mL) were dissolved in 1,4-dioxane (15 mL), the reaction was stirred at 40 °C for 1 h. And then the solution was concentrated to remove the solvent, dry HCl salt was obtained. On the other hand, compound **3-4** (953 mg, 5.0 mmol) and HATU (2.9 g, 7.5 mmol) were dissolved in DCM (40 mL) under $N_2$ atmosphere, the reaction was stirred at 25 °C for 0.5 h. Then DIPEA (1.9 mL, 11.0 mmol), and HCl salt were added to the reaction, the reaction mixture was stirred for 10 h at 25 °C. Then DCM (50 mL) and 1N HCl aqueous solution (10 mL) were added to the reaction. After separation, the organic phase was washed with water (10 mL) and saturated brine (10 mL), dried over anhydrous sodium sulphate, concentrated under vacuum and purified by column chromatography to give compound **5-3** as white solid (1.5 g, yield: 57.7%).

**(*S*)-*N*-((*S*)-1-((3*R*,5'*S*)-5'-cyano-2-oxospiro[indoline-3,3'-pyrrolidin]-1'-yl)-3-cyclopropyl-1-oxopropan-2-yl)-2-(2,2-difluoroacetamido)-*N*-methylpropanamide (5).** Compound **5-3** (400 mg, 0.76 mmol) and 4 *N* HCl in 1,4-dioxane (2 mL) were dissolved in 1,4-dioxane (2 mL), the reaction was stirred at 25 °C for 1 h. Then the reaction was concentrated to remove the solvent and $Et_3N$ (0.37 mL, 2.7 mmol), difluoroacetic anhydride (0.21 mL, 1.7 mmol), DCM (4 mL) were added to the reaction at 0 °C. The reaction was allowed to warm to 25 °C and stirred for 10 h. Then DCM (10 mL) and 1*N* HCl aqueous solution (3 mL) were added to the reaction. After separation, the organic phase was washed with saturated brine (5 mL), dried over anhydrous sodium sulphate, evaporated in vacuum and purified by column chromatography to give **5** as white solid (120 mg, yield: 32%). $^1$H NMR (400 MHz, DMSO-$d_6$) δ 10.71 (s, 1H), 8.94 (d, J = 7.4 Hz, 1H), 7.24 (t, J = 7.6 Hz, 1H), 7.04 (d, J = 7.4 Hz, 1H), 6.98 (t, J = 7.5 Hz, 1H), 6.89 (d, J = 7.8 Hz, 1H), 6.14 (t, J = 53.6 Hz, 1H), 5.22 (t, J = 7.4 Hz, 1H), 5.15 (t, J = 7.9 Hz, 1H), 4.59 (t, J = 7.0 Hz, 1H), 3.82–3.64 (m, 2H), 2.93 (s, 3H), 2.64 (dd, J = 13.3, 8.7 Hz, 1H), 2.49–2.42 (m, 1H), 1.60 (hept, J = 7.4 Hz, 2H), 0.79 (d, J = 6.9 Hz, 3H), 0.56 (dt, J = 13.7, 7.5 Hz, 1H), 0.33 (ddp, J = 22.2, 9.2, 4.9, 4.1 Hz, 2H), 0.06 (tt, J = 13.8, 7.0 Hz, 2H). $^{13}$C NMR (151 MHz, Chloroform-*d*) δ 175.75, 172.01, 170.18, 139.78, 131.05, 129.58, 123.23, 122.31, 116.77, 110.75, 108.19, 55.10, 54.74, 52.77, 46.04, 45.65, 39.12, 34.17, 30.90, 17.62, 7.44, 4.77, 4.45. ESI-HRMS Calcd for $C_{24}H_{28}F_2N_5O_4$ $[M+H]^+$: 488.2104, found 488.2089.

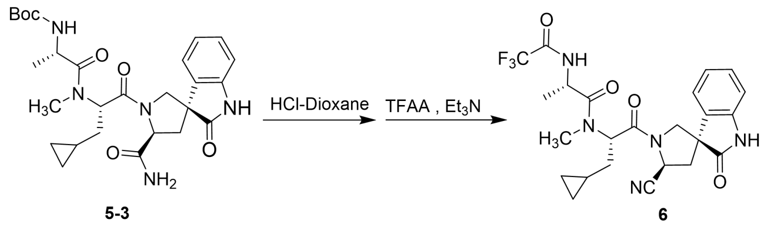


**Figure S6** The synthesis path of **6**.

**(*S*)-*N*-((*S*)-1-((3*R*,5'*S*)-5'-cyano-2-oxospiro[indoline-3,3'-pyrrolidin]-1'-yl)-3-cyclopropyl-1-oxopropan-2-yl)-*N*-methyl-2-(2,2,2-trifluoroacetamido)propenamide (6).** Compound **5-3** (400 mg, 0.76 mmol) and 4 *N* HCl in 1,4-dioxane (2 mL) were dissolved in 1,4-dioxane (2 mL), the reaction was stirred at 40 °C for 1 h.Then the reaction was concentrated to remove the solvent and $Et_3N$ (0.4 mL, 2.7 mmol), trifluoroacetic anhydride (0.2 mg, 1.67 mmol), DCM (4 mL) were added to the reaction at 0 °C. The reaction was allowed to warm to 25 °C and stirred for 10 h. Then DCM (5 mL) and 1*N* HCl aqueous solution (1 mL) were added to the reaction. After separation, the organic phase was washed with water (2 mL) and saturated brine (2 mL), dried over anhydrous sodium sulphate, concentrated under vacuum and purified by column chromatography to give 6 as white solid (150 mg, yield: 40.0%).$^1$H NMR (400 MHz, DMSO-$d_6$) δ 10.71 (s, 1H), 9.56 (d, J = 7.0 Hz, 1H), 7.24 (td, J = 7.7, 1.4 Hz, 1H), 7.05 (dd, J = 7.5, 1.3 Hz, 1H), 6.98 (td, J = 7.5, 1.0 Hz, 1H), 6.89 (d, J = 7.8 Hz, 1H), 5.23 (dd, J = 8.1, 6.8 Hz, 1H), 5.14 (dd, J = 8.7, 7.1 Hz, 1H), 4.59 (p, J = 7.0 Hz, 1H), 3.85-3.60 (m, 2H), 2.93 (s, 3H), 2.65 (dd, J = 13.2, 8.7 Hz, 1H), 2.53-2.42 (m, 1H), 1.61 (td, J = 7.2, 3.1 Hz, 2H), 0.83 (d, J = 7.0 Hz, 3H), 0.54 (dtd, J = 12.3, 7.3, 5.0 Hz, 1H), 0.41-0.19 (m, 2H), 0.11-0.01 (m, 2H). $^{13}$C NMR (151 MHz, Chloroform-*d*) δ 175.87, 171.61, 170.11, 139.82, 131.00, 129.62, 123.23, 122.30, 116.74, 114.64, 110.81, 55.24, 54.75, 52.80, 46.36, 46.07, 39.09, 34.16, 30.93, 17.41, 7.44, 4.79, 4.47. ESI-HRMS Calcd for $C_{24}H_{25}F_3N_5O_4$ $[M-H]^-$: 504.1859, found 504.187.

**Materials and methods**

All commercially available chemicals and solvents were directly used without further purification. All reactions were monitored by thin layer chromatography (TLC) on silica gel plates (GF-254). Molecular mass was determined on a mass spectrometry (Waters, VION IMS QTOF). High-resolution mass spectra (HRMS) were measured on Sciex ZenoTOF™ 7600 System. 1H NMR and 13C NMR spectra were recorded using a Bruker 400 MHz or 600 MHz spectrometer with tetramethylsilane as an internal standard.
